# Pan-cancer silencer transcription atlas reveals leukemia silencer hijacking FOXP1 to enhance *MYC* oncogene

**DOI:** 10.64898/2026.01.08.698306

**Authors:** Mengbiao Guo, Liutao Chen, Shengcheng Deng, Jingkai Zhang, Zhijie Hu, Weiwen Wang, Zhou Songyang, Yuanyan Xiong

## Abstract

Numerous non-coding regulatory elements have been identified in the human genome. However, our understanding of silencers remains largely incomplete. Here, we proposed the concept of silencer transcriptional activity (STA) to investigated silencers from a new perspective using >20,000 transcriptomes. We demonstrated that lower STAs predicted better patient outcomes in most cancers, and a subset of STAs distinguished tumors from noncancerous tissues for most cancer types. Importantly, we discovered a silencer-linked RNA (slRNA) of a leukemia cell-specific silencer (LeSSi). Knocking down (KD) of the slRNA significantly decreased leukemia cell viability and migration and increased cellular apoptosis. Furthermore, the direct link and functional similarity between LeSSi and slRNA were established by CRISPR/Cas9 knocking out (KO) of LeSSi. Mechanistically, the slRNA hijacked FOXP1, which functions here both as a known transcription factor (TF), and more importantly, as a novel RNA binding protein (RBP), thereby preventing FOXP1 from binding to the promoter of proto-oncogene *MYC* and subsequently promoting *MYC* transcription. Collectively, our work provides insights into the functions of silencers in cancer and identified a slRNA hijacking FOXP1 to enhance *MYC* expression in leukemia cells.

## Introduction

Silencers were genomic elements repressing target gene transcription, in contrast to enhancers which enhances gene transcription [1,2]. Early silencer research dates back to decades ago, such as the intronic silencer regulating *CD4* gene expression during T-cell development [3,4], and silencers in hypermethylated regions of *Igf2* which resulted in imprinting loss of the *H19* gene [5]. Recently, two studies have reported genome-wide silencer elements in human [6] and mouse [7]. Pang and Snyder devised an elegant screen system to examine human silencers in human open chromatin regions, using K562 (chronic myelogenous leukemia, CML) and HepG2 (hepatocellular carcinoma) cells [6]. Ngan et al. uncovered mouse silencers by studying chromatin interactions induced by polycomb repressive complex 2 (PRC2) in mouse embryonic stem cells (mESCs) [7]. Both studies found that many silencers may repress their target genes through long-range chromatin interactions.

Interestingly, some silencers may transit to enhancers dynamically during development, and 25% of PRC2-bound silencers in mouse embryonic stem cells may follow this pattern during differentiation [7,8]. Moreover, the Encyclopedia of DNA Elements (ENCODE) project suggested that most of the human genome (>90%) can be transcribed [9]. Inspired by transcription within enhancers [10] and the observation that silencers are enriched in weakly transcribed regions as indicated by epigenetic modifications [6,7], we reasoned that many silencers may have been covered by RNA sequencing (RNA-seq) and their RNA coverage may have biological meaning, which we termed as transcriptional activities of silencers (STAs). Of note, STAs may not only be resulted from stable transcriptional products of silencer host or surrounding genes, but also can be from unknown (unannotated) non-coding RNAs.

We hypothesized that high levels of STAs would affect silencer functions by preventing transcriptional factors or epigenetic factors from binding to silencers to function properly (**Figure 1A**). Moreover, long noncoding RNAs (lncRNAs) are transcribed pervasively across the human genome, however, functions of most lncRNAs are unknown [11–14]. Because many lncRNAs act as transcriptional repressors [15,16], differential STAs in diseases may help uncover disease-relevant lncRNAs overlapping silencers (termed silencer-linked RNAs, or slRNAs), and these slRNAs may have functions similar to silencers.

**Figure 1.** Overview of silencer transcriptional activities across cancer types. (**A**) The two concepts proposed in this study, STA and silencer-linked RNA (slRNA), and their hypothesized functions. High levels of STAs may block binding of transcription factors or epigenetic factors to silencers. slRNAs were identified by overlapping long non-coding RNAs (lncRNAs) with silencers showing interesting patterns of STAs. (**B**) t-distributed Stochastic Neighbor Embedding (tSNE) clustering of cancer samples across cancer types (indicated by different colors). (**C-D**) Distribution of significant Spearman’s correlation coefficients (mostly >0) between STA and DNA accessibility by assay for transposase-accessible chromatin with sequencing (ATAC-seq) (**C**), and one example of correlation between STA of a silencer (10kb downstream of *KDM4B*) and *KDM4B* accessibility in breast cancer was shown in (**D**). (**E**) Significant differences of silencer effects (fold changes over background) among silencer groups, with silencers located in promoter regions showing the smallest effect. The fold change information for each silencer was derived by comparing signals from cells treated with or without inserting the silencer fragment and was downloaded from Supplementary Table 1 of the published study (Pang and Snyder, Nature Genetics, 2020). (**F**) Overall STA difference between tumor and normal samples in each cancer type. STA: silencer transcriptional activities, ACC: Adrenocortical carcinoma, BLCA: Bladder urothelial Carcinoma, BRCA: Breast invasive carcinoma, CESC: Cervical squamous cell carcinoma and endocervical adenocarcinoma, CHOL: Cholangiocarcinoma, COAD: Colon adenocarcinoma, DLBC: Lymphoid neoplasm diffuse large B-cell lymphoma, ESCA: Esophageal carcinoma, GBM: Glioblastoma multiforme, HNSC: Head and neck squamous cell carcinoma, KICH: Kidney chromophobe, KIRC: Kidney renal clear cell carcinoma, KIRP: Kidney renal papillary cell carcinoma, LAML: Acute myeloid leukemia, LGG: Brain lower grade glioma, LIHC: Liver hepatocellular carcinoma, LUAD: Lung adenocarcinoma, LUSC: Lung squamous cell carcinoma, MESO: Mesothelioma, OV: Ovarian serous cystadenocarcinoma, PAAD: Pancreatic adenocarcinoma, PCPG: Pheochromocytoma and paraganglioma, PRAD: Prostate adenocarcinoma, READ: Rectum adenocarcinoma, SARC: Sarcoma, SKCM: Skin Cutaneous melanoma, STAD: Stomach adenocarcinoma, TGCT: Testicular germ cell tumors, THCA: Thyroid carcinoma, THYM: Thymoma, UCEC: Uterine corpus endometrial carcinoma, UCS: Uterine carcinosarcoma, UVM: Uveal melanoma.

Here, to test our hypotheses in a broad range of human tissue types and to obtain novel insights, we investigated STAs in a huge volume of RNA-seq samples (∼20,000), including both cancer and normal samples. Most significant STAs associations with DNA accessibility as reported in assay for transposase-accessible chromatin followed by sequencing (ATAC-seq) data were positive. We showed that STA levels were cancer-type specific, were generally upregulated across cancer types, and were prognostic for multiple cancer types. Moreover, a machine-learning model based on STAs of top 100 silencers can almost perfectly classify tumors and adjacent non-cancerous samples with high specificity and sensitivity across cancer types. Chromatin interaction analysis with paired-end tag sequencing (ChIA-PET) data revealed important long-range putative targets, for example, *ATM* and *EZH2*, of about half of these top silencers. Importantly, we identified a slRNA of a silencer (LeSSi) with STA specifically detected in K562 cells. The slRNA was capable of enhancing *MYC* gene expression by hijacking the FOXP1 protein, which otherwise bound to the *MYC* promoter to repress its transcription. Collectively, our work introduced two new concepts (STA and slRNA) to study silencers and revealed the mechanism of a functional slRNA in leukemia cells.

## Results

### Pan-cancer characterization of STAs

First of all, we found enrichment of transcription signals, as indicated by cap analysis of gene expression (CAGE)-seq peaks (**Figure S1A**), as well as transcription-friendly epigenetic context, as indicated by ATAC-seq signals (Figure S1B), surrounding silencer regions. Silencers were also found to be enriched in weakly transcribed regions, according to previous research [6]. These results confirmed that our STA hypothesis for silencers (**Figure 1A**) is sound. Then, for >11,000 RNA-seq samples across 33 cancer types from The Cancer Genome Atlas (TCGA), we calculated normalized STAs (defined as active if STAs >= 5) by examining read coverage in 2,661 experimentally-defined silencer regions in K562 cells [6] (see **Methods**). We then performed t-SNE analysis of STAs and found that most cancer samples clustered by tissue types (Figure 1B), indicating that STAs represented a biological dimension of interest and there exist probably cancer-specific STAs.

We further performed two types of analyses to support our STA hypothesis. First, it is expected that active polymerase transcription in the silencer region (i.e. high STA) should have positive correlation with high chromatin accessibility of the region. Indeed, we observed 635 pairs of significant correlation between STAs and previously reported ATAC-seq peak accessibility [17], and most pairs (97.3%) showed positive correlations (Figure 1C, **Table S2**). The best positive correlation, observed in breast cancer (BRCA), of a silencer, located 10kb downstream of *KDM4B*, was shown as an example (Figure 1D). Interestingly, potential targets of the associated ATAC-seq for this silencer included *KDM4B*, *UHRF1*, *DPP9*, *TICAM1*, *SAFB2*, and *SAFB1*. Of note, abnormal activities of *KDM4B* (histone demethylase for H3K9me3) and *SAFB1/2* (scaffold attachment factor B associated with estrogen receptor regulation) were reported in breast cancer [14,18]. Second, we compared silencer signal strength (the fold changes of silencer over background control reported by [6]) between the three main categories of silencers, located in promoter, intron, or distal intergenic regions, respectively. Interestingly, we found that promoter silencers, presumably more openly accessible than introns and intergenic regions, generally showed lower silencing effects than intronic and intergenic silencers (Figure 1E). Third, consistent with our STA hypothesis, we observed a trend of negative association between STAs and the silencer signal strength (Figure S1C).

### STAs were upregulated and prognostic across cancer types

Next, we examined overall STA difference between tumor and matched normal samples. Most tumor samples showed higher STAs than tumor adjacent normal samples, especially for CHOL, ESCA, HNSC, KIRC, LIHC, and STAD, except for KICH and THCA (Figure 1F, **Figure S2**). Consistently, we observed that lower STAs were associated with better patient survival in most cancer types with nominal significance (*P* < 0.05, **Figure S3**), including significance in KIRC (*P* =1.1e-4) and LGG (*P* = 2.8e-3) after multiple testing correction. Interestingly, HNSC prognosis seems different compared to the other cancer types, which may be due to prevalent HPV infection in HNSC [14,19].

### The majority of prognostic STAs outperformed their nearest genes

We then identified individual prognostic STAs and compared them with their nearest genes (or host genes, if silencers located in exons, introns, or untranslated regions [UTR]). A total of 188 STAs were prognostic (FDR < 0.1) across cancers, but only 83 (44.1%) of the nearest/host genes with expression levels prognostic for patient outcomes after multiple correction (FDR < 0.1) (**Figure 2A**, **Table S3**). These 83 nearest/host genes were most likely true targets of corresponding silencers. Consistent with the general upregulation of STAs in tumors, higher levels of most prognostic STAs predicted worse outcomes of patients; two examples, silencers with nearest/host genes *DHX32* and *CDR1*, respectively, were shown (Figure 2B-C). More importantly, expression levels of all these prognostic nearest genes resulted in concordant risk classification (higher values predicted either risk or protective effects) when compared with their corresponding STAs. Two significant silencers and their nearest/host genes, *HOXA3* and *LRP5*, respectively, were also shown as examples (**Figure S4**). These findings added clinical support to the utility of cancer STAs.

**Figure 2.** Prognostic STAs and nearest/host genes. (**A**) Comparison significance (FDR) between STAs (x-axis) and their nearest/host genes (y-axis). Nearest/host gene names of the top ten STAs were labelled. A diagonal line of y = x and a horizontal line where GeenFDR = 0.1 were shown. (**B-C**) Two examples of prognostic STAs (left panel), chr10:125853371-125853550 (**B**) and chrX:140783169-140784666 (**C**), with non-prognostic nearest/host genes (DHX32 and CDR1, respectively) (right panel) in LGG. Survival patient groups were defined by using 33% and 67% quantiles ranked by STA levels. Log-rank test *P*-values were shown. Yellow color indicates the patient group with low gene expression and green for high gene expression.

### Top STAs associated with protein kinases were highly predictive of pan-cancer tumors versus non-tumors

To further explore the clinical significance of individual STAs, we built a random forest-based pan-cancer machine learning model to examine whether STA features can separate tumors from adjacent normal samples across cancer types (**Figure S5A**). After obtaining STA importance in predicting tumors, we found that the top 100 STAs (**Table S4**) achieved the best performance, with an overall sensitivity of 0.992 and specificity of 0.857 (for cancers with matched normal samples), across cancers (**Figure 3A**, values of performance shown in **Table S5**). The moderate specificity was probably due to both the small number of normal samples and large heterogeneity (unbalance of tumor and normal samples) across cancers. Remarkably, the model achieved 100% specificity for CHOL, LIHC, LUAD, LUSC, and READ, and close to or above 90% for BRCA, COAD, HNSC, KICH, KIRC, STAD, and THCA. Next, we examined these top 100 silencers, most of which were located in promoter regions. Their nearby genes were closely related to cancer and inflammation, such as *PSMB4*, *LATS1*, *IGF2BP2*, *CD40*, *COQ8A*, *PRDM2*, and *RIPK3* (Figure 3B). The most enriched functions of genes close to these silencers were protein serine/threonine kinase activity and ubiquitin-like protein ligase binding (Figure S5B). Interestingly, the majority of these top silencer-related genes were prognostic for patients across multiple cancer types and higher expression levels were associated with higher cancer risk (**Figure S6**), as examined by using the GEPIA2 webserver [20]. Of note, *IGF2BP2* was recently reported to be a potent oncogene in cancer, by binding to N6-methyladenosine (m6A) modifications on diverse mRNAs, including MYC transcripts, to stabilize them [21].

**Figure 3.** Top STA features identified by machine learning were highly predictive of tumor samples versus adjacent normal samples. (**A**) Model performances based on different sets of features were shown. The best K = 100 STAs was chosen (indicated by an arrow) for further analysis, according to testing sensitivities and specificities. (**B**) Nearby genes of the top 200 silencers based on STA importance from machine learning. Colors represent different genomic regions (mostly promoter regions, pink). Genes of interest for STAs were labelled. (**C-D**) Visualization of Pol2 Chromatin interaction analysis with paired-end tag sequencing (ChIA-PET) silencer-promoter interactions linking silencers to putative long-range target genes, ATM/CUL5 (**C**) and EZH2 (**D**). (**E**) Spearman’s correlation between silencer STA (cyan color) or silencer-proximal gene expression (orange color) and ATAC-seq target gene ATM across cancer types (x-axis). Significance indicated: * *P* <= 0.05, ** <= 0.01, *** <= 0.001, or a dot for non-significance.

We further examined these top silencers by investigating their putative long-range target genes using Pol2 ChIA-PET data from K562 cells [22], in addition to their proximal genes shown above. Among the top 100 silencers, 42 overlapped with ChIA-PET interacting regions that targeting 78 genes, including *ATM, CUL5, E2F4, EZH2, H2BC11, H2BC12, ICAM1, ICAM5, MAVS, POU5F1, RPS2,* and *ZKSCAN3* (Table S4). Two silencer examples were shown. The first one was chr11-108121229-108121767 (hg38, termed slATM) targeting *ATM* (serine/threonine protein kinase playing critical roles in in DNA double strand breaks [DSB], together with *TP53* and *BRCA1*, and many other important cellular processes [23,24]) and *CUL5* (core component of E3 ubiquitin-protein ligase complexes mediating proteasomal degradation of target proteins [25]) (Figure 3C). The second one was chr7-149125749-149125922 (hg38, termed slEZH2) targeting *EZH2* (core component of the epigenetic complex PRC2 that regulates histone tail methylation [26] and is close associated with silencers [7]) (Figure 3D). Remarkably, slATM STA showed much more significant correlation with ATM expression than the slATM nearby gene (*ACAT1*) expression across most cancer types (Figure 3E), and similar results were observed for slATM and CUL5 (**Figure S7A**). In contrast, slEZH2 STA showed stronger correlation with EZH2 expression than the slEZH2 nearby gene (*ZNF425*) expression only in several cancer types (Figure S7B). The pan-cancer positive correlation between STA and target gene expression was consistent with our STA hypothesis.

### STA profiles revealed a leukemia cell-specific silencer-linked RNA

We further computed STAs in 102 K562/CML cell line samples and >9,000 normal samples across 30 tissues from the Genotype-Tissue Expression project (GTEx) to identify potentially important K562-specific silencers. We then summarized the transcriptional activities by calculating the fraction of samples with active (>= 5 reads) silencer transcription in each tissue or cancer for each silencer. We observed that STAs were highly consistent across both cancer types and tissue types, and they were detected mostly in a subset of silencers located in promoters, followed by exons, introns, UTRs, and intergenic regions (**Figure S8**). Next, we focused on the top 100 silencers as ranked by the proportion of active STAs in K562 (**Figure 4A**). Interestingly, several STAs were exclusively detected in K562/CML, including an intergenic silencer (ranked 50^th^, chr18:77918019-77918253, named LeSSi) overlapping an exon of the lncRNA T164574 (a potential slRNA, renamed to LeSSiR for better annotation), which was annotated in the MiTranscriptome database [13] and was included in our previous lncRNA collection [14] (Figure 4B). To test the fidelity of our analysis, we then validated the transcriptional product of the slRNA (LeSSiR) using RT-PCR (Figure 4C) and Sanger sequencing (Figure 4D, **Figure S9**).

**Figure 4.** STAs across 21,000 samples reveal a K562/CML-specific slRNA validated by CRISPR/Cas9-based silencer knockout. (**A**) The top 100 STAs ranked by K562/CML STAs. CML: chronic myeloid leukemia. Rows are cancer types or tissue types and columns are silencers. Tissue types and genomic regions for silencers were shown. K562, TCGA-LAML (acute myeloid leukemia), and a K562-specific silencer (LeSSi) was indicated by blue arrows at the bottom and a circle at the top row for K562 samples. (**B**) Overview of genomic regions around the silencer-linked RNA (slRNA) (T164574, renamed to LeSSiR) in UCSC Genome Browser. Tracks from top to bottom: genome scale, slRNA, and silencer. (**C**) The gel image showing reverse transcription-polymerase chain reaction (RT-PCR) products of the slRNA. Two products were observed for T164574 (LeSSiR) and the red arrow indicated the long isoform used for further experiments. (**D**) Screenshots of Sanger sequencing results for the PCR product shown in (**C**). (**E**) (top panel) Schematic showing silencer DNA deletion (knockout, KO) by clustered regularly interspaced short palindromic repeats (CRISPR)/Cas9 with paired guide-RNA to introduce large fragment deletions. (bottom panel) Silencer KO led to the downregulation of the slRNA T164574 (LeSSiR) expression. T: T164574, LeSSi: Leukemia-specific silencer element, LeSSiR: LeSSi RNA. Two-sided t-test. * P <= 0.05, ** <= 0.01, *** <= 0.001.

To further establish the direct link between silencer DNA and the slRNA, we deleted the silencer DNA fragment of the slRNA in K562 cells by using the CRISPR/Cas9 knockout (KO) system [27,28] for large genomic fragment deletions. We observed significantly reduced slRNA upon KO of the silencer (Figure 4E), which confirmed our hypothesis of the direct link between silencer DNA and the slRNA.

### Knockdown of the slRNA suppressed leukemia cells viability potentially mediated by FOXP1

To identify biological functions of the slRNA (LeSSiR), we used RNA interference (RNAi) to knock it down in K562 cells (**Figure 5A**). Cell viability assays showed that slRNA knockdown (KD) suppressed leukemia cells (Figure 5B). Transwell assay results further showed that slRNA KD reduced tumor ability of cell migration (first step of tumor metastasis [29]) (Figure 5C). Moreover, flow cytometry results showed that slRNA KD increased cell apoptosis (Figure 5D).

**Figure 5.** Cancer-associated biological functions of slRNAs. (**A**) Knockdown (KD) effects of two siRNAs for the slRNA. T164574 (LeSSiR) expression was reduced after treatment of siRNAs. (**B-D**) slRNA KD suppressed K562 cells viability (**B**) and cell migration (**C**) and increased cellular apoptosis (**D**). Two-sided t-test. * *P* <= 0.05, ** <= 0.01, *** <= 0.001.

We then performed RNA sequencing of slRNA-KD and control samples. RNA-seq results showed that slRNA KD significantly upregulated some genes (differentially expressed genes, DEG), including the p53-inhibitor gene *MDM4*, and downregulated some genes, including the potent proto-oncogene *MYC* (**Figure 6A**). We further examined expression changes of the representative genes (*MYC* and *MDM4*) in silencer KO cells established above by CRISPR/Cas9. We observed consistent results (significant down-regulation of *MYC* and up-regulation of *MDM4*) between slRNA KD and silencer DNA KO (Figure 6B).

**Figure 6.** slRNA hijacked FOXP1 to prevent it from binding to the *MYC* promoter. (**A**) Differentially expressed genes (DEGs) after slRNA KD. Two top DEGs, *MYC* and *MDM4*, were labelled in blue. NS, not significant. X-axis for gene expression was truncated for better visualization. (**B**) Knocking out (KO) the silencer upregulated *FOXP1* and *MDM4* expression and downregulated *MYC* expression. T: T164574. (**C**) Significantly enriched transcription factors (TFs) in up- or down-regulated DEGs for slRNA KD. Red colors indicate FDR < 0.1. Top TFs were labelled and FOXP1 was indicated by an arrow. (**D**) Chromatin immunoprecipitation (ChIP)-qPCR experiments with two FOXP1 antibodies, #1 and #2, to examine enrichment of *MYC* promoter sequence in FOXP1-bound DNAs. (**E**) RNA Immuno-Precipitation (RIP) RT-PCR experiments showing that FOXP1 bound to the slRNA. (**F**) Additional FOXP1 KD in K562 cells with slRNA KD restored K562 cells viability. (**G**) The proposed model of slRNAs regulating FOXP1 and *MYC* by hijacking FOXP1 in K562 leukemia cells. (left panel) Under normal condition, slRNAs were not expressed and FOXP1 bound to the *MYC* promoter to repress *MYC* transcription. Silencers can also repress FOXP1 expression. (right panel) In leukemia cells, slRNAs were upregulated and interacted with FOXP1 to prevent it from binding to the MYC promoter, subsequently enhancing *MYC* expression and promoter cancer cell viability and migration but repress cell apoptosis. (**B, D, F**) Two-sided t-test. * *P* <= 0.05, ** <= 0.01, *** <= 0.001.

To identify the possible mechanism of the slRNA, we performed transcription regulator enrichment analysis [30] of the up- and down-regulated genes after slRNA KD. We found that the slRNA possibly interacted with the transcription factor (TF) FOXP1, which ranked top and was highly significant (*P* < 1e-7) among up-regulated genes (top 100) but less so among down-regulated ones (top 100) (Figure 6C). Interestingly, FOXP1 was up-regulated upon silencer KO (Figure 6B), which was not observed in slRNA KD and may reflect certain functional difference between slRNA and silencer DNA.

### slRNAs interacted with FOXP1 to relieve inhibition of MYC expression

FOXP1 was reported to affect *MYC* through different mechanisms. For example, FOXP1 can bind to the *MYC* promoter in response to TGF-β signaling and was required to repress MYC expression in T cells [31]. Proteomic profiling also showed that FOXP1 had protein-protein interaction (PPI) with MYC [32]. We hypothesized that FOXP1 probably regulated MYC expression in K562 cells by binding to the *MYC* promoter. To verify the possible mechanism, we first confirmed the role of FOXP1 in *MYC* expression regulation, and results showed that KD of FOXP1 (**Figure S11A**) increased *MYC* expression in K562 cells (Figure S11B). Three FOXP1 siRNAs showed consistent effects in repressing FOXP1 and enhancing *MYC* expression levels. To further test whether the slRNA (LeSSiR) interferes *MYC* expression by affecting FOXP1 binding to the *MYC* promoter region in K562 cells, we performed chromatin immunoprecipitation qPCR (ChIP-qPCR) experiments using FOXP1 antibodies followed by qPCR. Remarkably, all ChIP-qPCR results showed enrichment of *MYC* promoter sequences (more than four-fold) after slRNA KD (Figure 6D). Therefore, slRNAs may interact with FOXP1 to relieve inhibition of MYC expression.

Although FOXP1 acts as a TF that interacts with DNA, FOXP1 possibly can also act as an RNA binding protein (RBP) and interact with slRNAs, because many RBPs are able to interact with chromatins to exert their biological functions [33]. Consistently, we observed scattered FOXP1 binding motifs (TGTTT) [34] within the slRNA sequence (**File S1**). RPIseq (http://pridb.gdcb.iastate.edu/RPISeq) [35] also computationally predicted high probabilities (0.99) of FOXP1 binding to the slRNA. To further support our hypothesis, we used RIP-PCR to demonstrate that the slRNA indeed physically interacted with FOXP1 (Figure 6E).

To corroborate these results, cell phenotypic experiments have also shown that FOXP1 KD rescued K562 cells viability when the slRNA was knocked down (Figure 6F). Based on the above results, we proposed our slRNA model: Under normal condition, slRNAs were not expressed and FOXP1 bound to the *MYC* promoter to repress MYC transcription. Silencers can also repress FOXP1 expression (Figure 6G, left); In leukemia cells, slRNAs were upregulated and interacted with FOXP1 to prevent it from binding to the MYC promoter, subsequently enhancing *MYC* expression and promoter cancer cell viability and migration but repress cell apoptosis. Silencers also failed to repress FOXP1 expression (Figure 6G, right). Our model can also explain abnormally high levels of FOXP1 in AML samples among all cancer types (**Figure S12**).

## Discussion

In this work, we investigated silencers at the transcription level in a large number of RNA-seq samples across multiple cancer types and tissues. To the best of our knowledge, this is the first study proposing the concept of STAs and slRNAs, and providing evidence to support their biological functions and clinical significance. Importantly, we demonstrated that a slRNA (LeSSiR) interact with FOXP1 that functions as both a TF (binding to the oncogene *MYC* promoter; known) and an RNA binding protein (interacting with slRNAs; novel) and prevent FOXP1 from binding to the *MYC* promoter, which subsequently promotes *MYC* transcription in leukemia cells.

Despite the fact that silencers may be dual functional (silencer/enhancer) regulatory elements (DFRE), DFRE silencers have been recently estimated to be around 6% [8]. This is also consistent with our observations that (i) only 1% of all silencers used in this study overlapped with Functional Annotation of the Mammalian Genome (FANTOM) enhancers previously surveyed across TCGA cancer types [36]; (ii) most active STAs were consistently detected in a subset of silencers across cancer types and tissue types, similarly as in K562 cells in which all silencers were experimentally identified [6].

For patient prognosis prediction, although STAs outperform their nearest genes in number, in terms of the statistical power of prognosis, among the STA nearest genes with statistical significance, expression of many genes outperformed STAs, probably due to higher noise level in STAs than within gene expression. Moreover, interestingly, the STA outperformance tended to be cancer-specific (KIRC and LGG). The reason is unclear and need further investigation in the future.

Since the FOXP1-regulated *MYC* itself was enriched in TFs regulating DEGs of slRNA knockdown, many DEGs were probably regulated by the slRNA/FOXP1/MYC axis. We noticed that the Treg marker FOXP3, also enriched in TFs regulating those DEGs, can also suppress *MYC* transcription to reprogram Treg metabolism [37], which we speculate may also be related to slRNAs and warrants further research.

Collectively, we proposed new methods to investigate silencers, especially in leukemia cells. Our STA atlas datasets presented here provide a starting point for further exploration by the research community. Future research combining our analytic strategy with more experimentally verified and computationally predicted silencers [8,38] may further improve our understanding of silencer mechanisms in cancer and other diseases.

## Methods and Materials

### Calculating STA

For all RNA-seq samples, including TCGA, GTEx, and K562, we downloaded uniformly processed sequencing data (in BigWig format which contains only sequencing frequencies of each nucleotide in the reference human genome, for sharing data faster and without privacy problems, as compared to BAM files containing the sequenced nucleotides) from Recount2 [39]. Then we calculated for each sample the STAs of K562 silencers (experimentally-defined regions without expansion), which were mostly located in noncoding regions [6], by counting reads mapped to silencer regions by using the function ‘coverage_matrix’, similar to previous studies [36,40]. STAs of each sample were normalized by total sequencing coverage that considers both region size and library size, by using the ‘scale_counts’ function from the recount R package (https://bioconductor.org/packages/release/bioc/html/recount.html). The STA estimation is very similar to the calculation of gene expression, except different genomic regions (silencer element regions vs. gene body regions) were used to count the reads mapped to these regions. According to the Recount2 study [39], a normalized read count of five was used to determine if STA was detected (active) or not in a sample. We further examined the overlap between silencers and enhancers with expression detected in TCGA [36] and found only 27 silencers overlapped with enhancers.

### Survival analysis

Survival analysis was performed by using the R package TCGAbiolinks (v2.14.1). The patient groups were defined by separating samples based on 33% and 67% quantiles. After sorting samples by STAs (or gene expression levels) from the highest to the lowest. STAhigh (or GeneHigh) group was defined as first 33% samples and STAlow (or GeneLow) group as the last 33% samples. Pan-cancer TCGA gene expression levels in transcripts per million mapped reads (TPM) were downloaded from the UCSC Xena database (https://xenabrowser.net/datapages).

### Transcriptional context analysis of silencers and enhancers

CAGE-seq (cap analysis of gene expression) [41,42] data for K562 cells were downloaded from the FANTOM5 database (https://fantom.gsc.riken.jp/5). BigWig format files (genomic read coverage) of ATAC-seq data for K562 cells were downloaded from the ENCODE database [43] (https://www.encodeproject.org/atac-seq). All silencer regions experimentally defined previously were aligned (stacked) by their centers and were expanded to be of the same size of 2000 bp (upstream and downstream 1000 bp relative to the silencer region center). The 5’-end positions of CAGE-seq reads were extracted and mapped to the expanded silencer regions (2000bp). The number of reads mapped to the 2000bp silencer regions were summarized by 10bp tandem windows (or bins) and were normalized by the average number of reads of all 10bp windows in the 2000 bp regions. For ATAC-seq data, average values of base-wise read coverage from the BigWig files were calculated across the stacked 2000 bp regions. Experimentally verified enhancers by massively parallel reporter assay (MPRA) were downloaded from a previous study [44].

### Machine learning for tumor-normal classification by STAs

For each cancer type, samples were randomly separated into training set (70% samples) and testing set (30%). First, random forest (from ‘scikit-learn’) was applied to the training set to obtain all features importance using the leave-one-out cross-validation strategy. Then, the top K (K = 50, 100, 200, 500, 2661/all) important features were validated using the testing set. For each chosen K, the subset training matrix, with only those top K features, were used as the input for retraining as in the first step, and this time the subset test matrix, also with only those top K features, were used to evaluate the model accuracy. Finally, the best K was chosen for downstream analysis.

### ChIA-PET data analysis

Preprocessed Pol2 ChIA-PET interactions were downloaded from a previous study [22] from a public database (https://www.ncbi.nlm.nih.gov/geo/query/acc.cgi?acc=GSE33664). ChIA-PET is one of the most accurate experimental methods to identify long-range interaction between regulatory elements and gene promoters. Silencer regions were overlapped with interaction region pairs (promoter-promoter or promoter-other) of ChIA-PET data, and then target promoter regions of silencer-overlapping ChIA-PET interactions were extracted. Spearman correlation between STA and the corresponding predicted silencer target gene were further conducted, which showed expected positive correlation, to support our STA hypothesis and accurate silencer target gene prediction based on ChIA-PET. Visualization of Pol2 ChIA-PET interactions for ATM/CUL5 and EZH2 were created by the 3D Genome Browser (http://3dgenome.fsm.northwestern.edu) [45].

### Pan-cancer ATAC-seq data analysis

Preprocessed pan-cancer ATAC-seq data from TCGA were downloaded from a previous study [17]. The nearest ATAC-seq peak was matched to each silencer. Spearman correlation and P-values between silencer STA and ATAC-seq peak accessibility were calculated by the cor.test function in R language in each cancer type. Association P-values were adjusted by FDR and FDR < 0.1 was considered significant.

### Cell culture and siRNA transfection

K562 cells were maintained in Iscove’s Modified Dulbecco’s Media (IMDM) (Gibco, 31980030) supplemented with 10% fetal bovine serum (Gibco, 10099141C) and 100 units/ml penicillin and 100 µg/ml streptomycin (Gibco, 15140122). The siRNA oligos were designed and synthesized by RiboBio (Guangzhou), and the siRNA target sequences were as follows: control (RIBOBIO CO., LTD); T164574-siRNA1: GCCAGATATTCCTTAATCA, T164574-siRNA2: GACACCACAAGGGATAATA; FOXP1-siRNA1: CTCAGTCCACACTCCCAAA, FOXP1-siRNA2: CCACAGAGCTTACCTCATA, FOXP1-siRNA3: CTGGTTCACACGAATGTTT. Cells at ∼40% density were transfected with siRNAs at a final concentration of 50 nM using Lipofectamine RNAiMAX Reagent (Invitrogen, 13778150) according to the manufacturer’s recommendations.

### RNA sample isolation and RNA sequencing analysis

K562 cells were transfected with siRNA at a final concentration of 50 nM and harvested with trypsin after treatment for 48h. Total RNA was extracted with TRIzol (Invitrogen, 10296010) according to the manufacturer’s protocol. RNA yield and quality were determined by UV absorption on a NanoDrop 1000 spectrophotometer and fragment size was analyzed using the RNA 6000 Nano assay (Agilent Technologies, Santa Clara, CA) run on the 2100 Bioanalyzer. Total RNA was used with the TruSeq RNA Library Preparation Kit v2, Set A (Illumina, San Diego, CA). The final pooled libraries were quantified and sequenced with Illumina NovaSeq 6000 Sequencing System. For quality control, raw reads were filtered by seqtk (v1.3, https://github.com/lh3/seqtk) to obtain clean reads as follows: (i) removing adapter sequences; (ii) removing 3’ bases with sequencing quality < 20; (iii) removing reads shorter than 25nt; (iv) removing ribosome RNA reads. Clean reads were mapped to the GRCh38 reference genome using HISAT2 (v2.0.4) [46]. Gene expression levels (read counts) were quantified with StringTie (v1.3.0) [47] and subjected to differential genes identification with edgeR [48] by |log2 fold change| ≥ 2 and adjusted P value ≤ 0.05). Gene ontology enrichment was performed using clusterProfiler (v3.14.3).

### Cellular viability, apoptosis and migration assays

Cell viability was analyzed by Cell Counting Kit-8 (abcam, ab228554) according to the manufacturer’s protocols. Briefly, Cells were seeded and cultured at a density of 1×10^4^ per well in 250 μL of medium into 48-well microplates (Thermo Scientific, 174898). Cells were transfected with 50 nM siRNA using Lipofectamine RNAiMAX Reagent (Invitrogen, 13778150). 10 μL of CCK-8 reagent was added to each well and then cultured for 2 hours after treatment for 48h. The absorbance was measured at 450 nm using a GloMax Discover System (Promega, GM3000) according to the manufacturer’s recommendations. The viability of cells was analyzed by the absorbance. Three biological repeats were performed for all experiments.

Cellular apoptosis was tested by Annexin V, FITC Apoptosis Detection Kit (Dojindo) and PE, 7-AAD Apoptosis Detection Kit (Dojindo). Briefly, 60 hours before apoptosis detection, camptothecin (8 μM) was added to each well of 24-well culture plates (2×10^5^ cells/well) for inducing apoptosis. Cells were harvested stained using the Apoptosis Detection Kits, then analyzed by flow cytometry as instructions of the kits.

Cellular migration was evaluated by Cell Migration/Chemotaxis Assay Kit (abcam, ab235696) according to the manufacturer’s instructions. Briefly, 5×10^4^ K562 cells were seeded in the upper chambers of each transwell with 100 μl culture medium without FBS, and 600 μl culture medium containing 10% FBS was added to the lower chambers. 48 hours later, cells were stained with cell dye, and readed the fluorescence at Ex/Em = 530/590 nm. Calculate the number of cells invaded using the equation of the straight line obtained from Standard Curve.

### Genes clone and RT-qPCR

Total RNA was extracted with TRIzol (Invitrogen, 10296010) according to the manufacturer’s recommendations. cDNA were reversed by SuperScript III Reverse Transcriptase with genes specific primer in Table S1, then, cDNA as template were amplified and clone into pCMV-Myc vector for sanger sequencing. cDNA was synthesized with PrimeScript™ RT reagent Kit (TAKARA, RR037A) for next qRT-PCR. Beta actin was used as an endogenous housekeeping gene for normalization. The relative mRNA level was appraised by 2^−ΔΔCt^ method. The primer pairs of the genes used for qRT-PCR were included in Table S1. Data were shown as relative gene expression.

### ChIP-qPCR

The ChIP protocol was performed according to our in house protocol. Briefly, K562 cells were transfected with siRNA and harvested after 48 hours. Cells were cross-linked for 10 minutes at room temperature by 1% formaldehyde, and quenched with 0.125 M glycine for 5 minute. Subsequently, the cross-linked cells were washed with TBSE buffer (20 mM Tris-HCl pH 7.5, 1 mM EDTA, 150 mM NaCl) and lysed with 0,1% SDS lysis buffer (50mM HEPES-KOH pH7.5, 150mM NaCl, 2mM EDTA, 1% Triton X-100, 0.1% Sodium deoxycholate, 0.1% SDS) for 15mins. Then, cell nuclear pellets were Centrifuged at 4000rpm for 15mins at 4℃ and further lysed by 1% SDS lysis buffer (50mM HEPES-KOH pH7.5, 150mM NaCl, 2mM EDTA, 1% Triton X-100, 0.1% Sodium deoxycholate, 1% SDS). The chromatin pellets were Centrifuged at 4000rpm for 15mins at 4℃, decant supernatant. The chromatin complexes were resuspended with Shearing Buffer (1mM EDTA, 10mM Tris-HCl pH 7.6, 0.1% SDS) and sonicated using covaris (M220) for 25 mins. Sonicated lysates were precleared with protein G Dynabeads (Invitrogen, 10003D) for 1 hour at 4°C and were incubated overnight at 4°C with magnetic beads bound with antibodies against FoXp1 (CST, D35D10) normal rabbit IgG (CST, 2729). The immunoprecipitations were washed 4 times and were eluted in elution buffer (50 mM Tris-HCl pH 8.0, 10 mM EDTA, 1% SDS) for the subsequent reversed cross-linked process by incubation with proteinase K (Thermo Fisher, 26160) at 55°C for 6 hours. The reversed cross-linked DNA was purified by Phenol–Chloroform–Isoamyl Alcohol (Thermo Fisher Scientific, 327110025) and ethanol precipitation. ChIP-qPCR assay was performed using TB Green Premix Ex Taq (TAKARA, RR420A). The primer sequences are listed in Table S1. The enrichment of specific genomic regions was calculated relative to the input DNA according to a previous report [49].

### Formaldehyde RNA Immuno-Precipitation (faRIP)

We perform faRIP as described previously [50]. One 10-cm dish K562 cells were seeded per RIP condition with 6×10^5^ cells/dish. Protein-RNA complexes were crosslinked in vivo by incubating cells with 10 mL of PBS-Formaldehyde (0.1%) and then halted it by quenching for 5 minutes at room temperature after adding glycine to a final concentration of 125 mM. Cells were centrifuged for 5 minutes at 500 g and then washed them twice in 4 °C PBS. 100 μL of protein G-Dynabeads® beads were prepared by initial washing with 3 × 1 mL RIP lysis buffer (50 mM Tris-HCl pH 7.5, 150 mM NaCl, 10% glycerol 1% NP-40, 0.1% SDS, and 0.5% sodium deoxycholate) before being blocked and loaded with the relevant antibody (2–10 μg diluted in 0.3 mL of RIP lysis buffer +1% BSA w/v final) for 1 hour at room temperature. Beads were washed with 3 × 1 mL RIP lysis buffer and left on ice. Each cell pellet was lysed in 400 μL RIP lysis buffer supplemented with 1 mM DTT, protease inhibitors (Sigma), 5 μL f RNase inhibitors (Ribosafe, Bioline) and 2 μL/mL of Turbo DNase (Ambion) Samples were sonicated using a Bioruptor (High, 5 × [30s-ON/30s-OFF]) and cleared by centrifugation (16100 x g, 10 min, 4 °C). 300 μL of each sample was incubated with Dynabeads for 2 hours at 4 °C and 30 μL of lysate was kept as input (10%). Following incubation, the beads were washed with 2 × 1 mL RIP lysis buffer, 2 × 1 mL high salt RIP lysis buffer (adjusted to 500 mM NaCl, 5 min each on ice), and 2 × 1 mL RIP lysis buffer. We re-suspended the frozen beads in 56 μl of RNase free water and added 33 μL of 3× reverse-crosslinking buffer (3× PBS (without Mg or Ca), 6 % N-lauroyl sarcosine, 30 mM EDTA, 15 mM DTT (add fresh)), and 10 μl of Proteinase K (Life Technologies, catalog #AM9516), and 1 μl of RNaseOUT to both the re-suspended beads and input sample. Finally, RNA complex are degradated and reverse-crosslinking for 1 hour at 42 °C, then another 1 hour at 55 °C. The RNA were extracted using TRIzol for next RT-qPCR assay.

### Silencer knockout by CRISPR/Cas9

Silencer knockout was performed using the CRISPR/Cas9 system. Paired sgRNAs were designed for a silencer using the sgRNA design website ChopChop (http://chopchop.cbu.uib.no/) and cloned into the PX458-GFP and PX459-dsRED gene editing vectors. Subsequently, the paired sgRNAs were co-transfected into K562 cells via electroporation. Double-fluorescent positive cells were sorted into single clones using flow cytometry and placed into 96-well plates. Individual clones were expanded and screened for homozygous silencer knockout by junction PCR amplification of the targeted genomic region, followed by Sanger sequencing to confirm the presence of deletion mutations. Clones with confirmed homozygous silencer knockout were used for subsequent phenotypic and functional analyses. T164574-slRNA-gRNA1: GGTGTATCGGAAAGCCCCTG, T164574-slRNA-gRNA2: GCTCAGCCAGATGTCAAGTAA; T164574-junction PCR-F: GCAGGCATGTTTAATGCTCTCTCAATCTATGAC, T164574-junction PCR-R:TCCAAAAAGCCAAATGGAGATTGGGGA

### Statistical analysis

Statistical analyses in this study were performed in R language (v3.6, https://www.r-project.org) or GraphPad 7. Statistical tests were indicated for each type analysis. Briefly, the comparison of STAs between tumor and normal samples was performed by two-sided Wilcoxon test. Two-sided *t*-test were performed for wetlab experiments with replicates. Significance of differential genes were calculated by the R package edgeR. Significance of TF enrichment in DEGs were provided by LISA webserver. Survival analysis significance was calculated by log-rank test in R. Adjusted P-values (FDR) were calculated by the R function p.adjust.

## Data availability

We deposited the raw RNA-seq data of lncRNA knockdown to SRA database (available upon acceptance). RNA-seq data of TCGA samples and K562/CML cells together with other samples of normal tissues from the Genotype-Tissue Expression project were obtained from Recount2 [39].

## CRediT author statement

**Mengbiao Guo**: Conceptualization, Data curation, Formal analysis, Methodology, Software, Visualization, Writing – original draft, Writing – review & editing, Funding acquisition. **Liutao Chen:** Methodology, Validation, Funding acquisition. **Shengcheng Deng:** Methodology, Validation. **Jingkai Zhang:** Formal analysis, Visualization. **Zhijie Hu:** Methodology, Validation, Funding acquisition. **Weiwen Wang:** Software, Formal analysis. **Zhou Songyang:** Writing – review & editing. **Yuanyan Xiong:** Resources, Methodology, Writing – review & editing, Funding acquisition, Supervision.

## Competing interest

The authors declare no competing interests.

## Acknowledgements

The work was supported by National Natural Science Foundation of China (NSFC) (Grant number 92249303 and 32000450), Guangdong Basic and Applied Basic Research Foundation, China (2021A1515110972 and 2020A1515010293), and Fundamental Research Funds for the Central Universities - Sun Yat-sen University, China (23ptpy62). The authors would like to thank TCGA and GTEx for making their research data publicly available.

## Supplementary Figure Legends

**Figure S1 Supporting evidence for transcriptional activities in silencer regions in K562 cells**

**(A)** A K562 cell CAGE-seq signal peak (y-axis) in the center (indicated by the position 0 of the x-axis) of silencer or enhancer regions. (**B**) A K562 cell ATAC-seq signal peak (y-axis) in the center of silencer or enhancer regions. Silencer or enhancer regions were expanded to include flanking 1000 bp for visualization of signal background. (**C**) Density plot with contour lines showing a trend of negative association between STAs and silencer signal strength in K562 cells.

**Figure S2 Paired testing of difference of average STAs between matched tumors and adjacent normal samples**

Two-sided Wilcoxon test. ns: nonsignificant, * *P* <= 0.05, ** <= 0.01, *** <= 0.001, **** <= 0.0001.

**Figure S3 Kaplan-Meier curves for analysis of average STAs in cancers with nominal significance of *P*-values**

Yellow color indicate the low-STA patient group and green for high-STA group. Log-rank test *P*-values were shown.

**Figure S4 Kaplan-Meier curves for survival analysis of expression levels of nearest/host genes associated with silencers**

(**A-B**) Two examples of prognostic STAs (left panel), chr7:27115516-27115731 (**A**, markedly fewer samples in STAhigh or GeneHigh group because of their low expression) and chr11:68386711-68386934 (**B**), with prognostic nearest/host genes (right panel) (HOXA3 and LRP5, respectively). Yellow color indicate the patient group with low gene expression and green for high gene expression. Log-rank test *P*-values were shown.

**Figure S5 Model design of the STA machine learning framework for sample classification (step 1 to 7)**

**(A)** STA features importance were first obtained by training a random forest model with 5-fold cross-validation in training samples (70%) across cancer types. Then, top features were extracted out from the full training matrix and full testing matrix to retrain the model by the subset of training features and test its performance on the subset of testing data. (**B**) Gene ontology (GO) enrichment of top 100 STA nearby genes.

**Figure S6 Most genes nearby the top 100 STAs were prognostic in multiple cancer types** Heatmap values are log10 hazard ratio (HR), and red or blue means higher expression for higher risk or lower risk, respectively.

**Figure S7 Correlation between STA or silencer nearby gene expression and expression levels of silencer targets based on ATAC-seq data**

(**A**) CUL5. (**B**) EZH2.

**Figure S8 The heatmap of all STAs ranked by K562/CML STAs**

Rows are cancer types or tissue types and columns are silencers. Tissue types and genomic regions for silencers were indicated. K562 was indicated by the red arrow.

**Figure S9 Example screenshots of Sanger sequencing and BLAST results for the RT-PCR product of slRNA T164574.**

**Figure S10 FOXP1 knockdown by siRNAs affected MYC expression**

**(A)** Three designed siRNAs targeting FOXP1, with gradually increasing knockdown (KD) effects on FOXP1. (**B**) Knocking down FOXP1 enhanced MYC expression, consistent with KD effects on FOXP1 for each FOXP1 siRNA. Two-sided t-test. *P < 0.05, ** <= 0.01.

**Figure S11 FOXP1 expression distribution across all cancer types of TCGA**

Created by using the GEPIA website (http://gepia2.cancer-pku.cn).

## Supplementary Table Legends

**Table S1** Primer sequences in qRT-PCR and ChIP-qPCR.

**Table S2** Significant associations between STAs and ATAC-seq accessibility across cancers.

**Table S3** List of prognostic STAs across cancers.

**Table S4** Information of the top 100 STAs of the pan-cancer machine learning model.

**Table S5** STA machine learning model performance across cancers.

## Supplementary Files

**File S1** FOXP1 binding motifs in slRNA T164574.

## Notes

### Competing Interest Statement

The authors have declared no competing interest.

