## Supplementary figures and images for "Pan-cancer silencer transcription atlas reveals leukemia silencer hijacking FOXP1 to enhance *MYC* oncogene"

### Supplemental Figures

Fig. S1

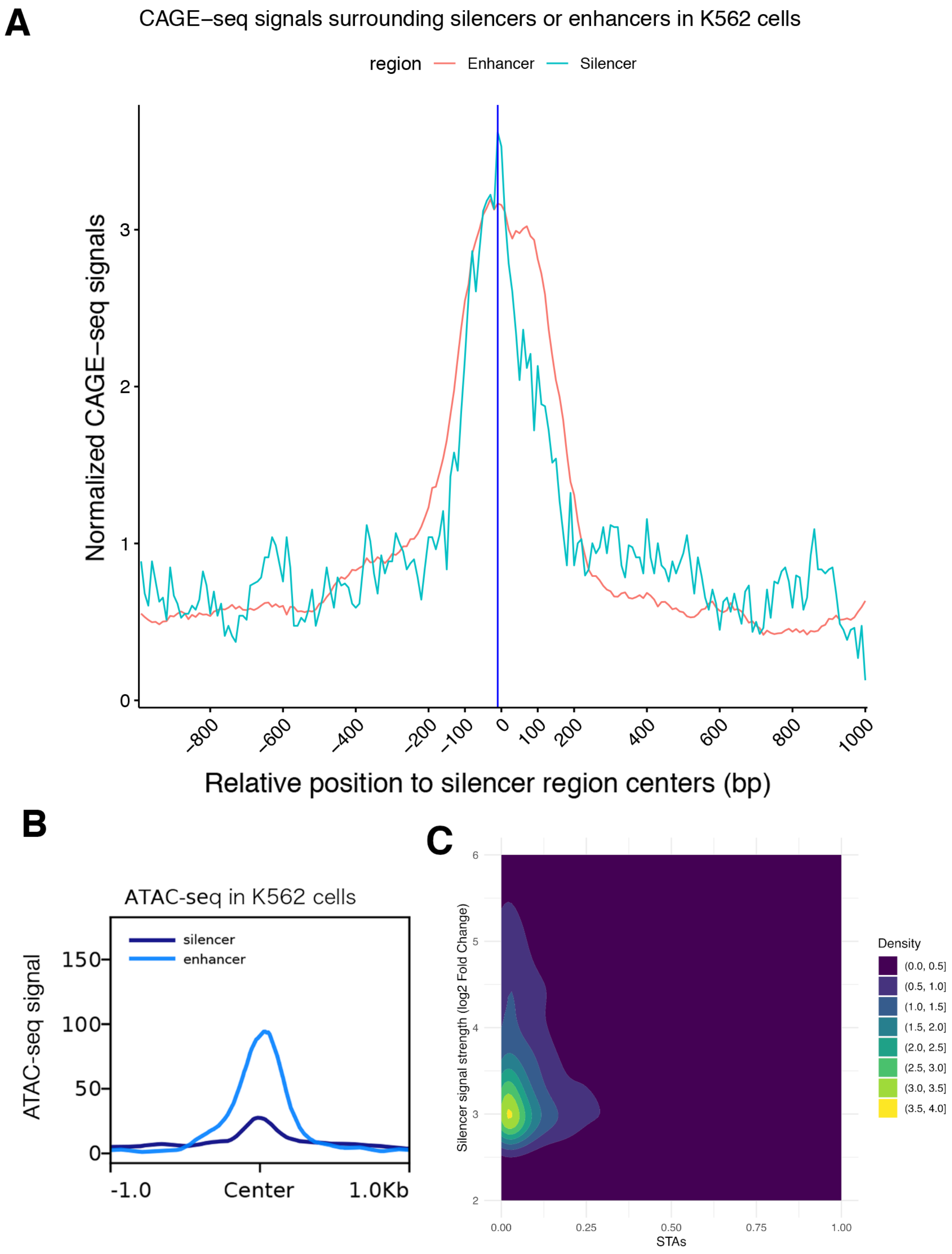


**Fig. S1.**

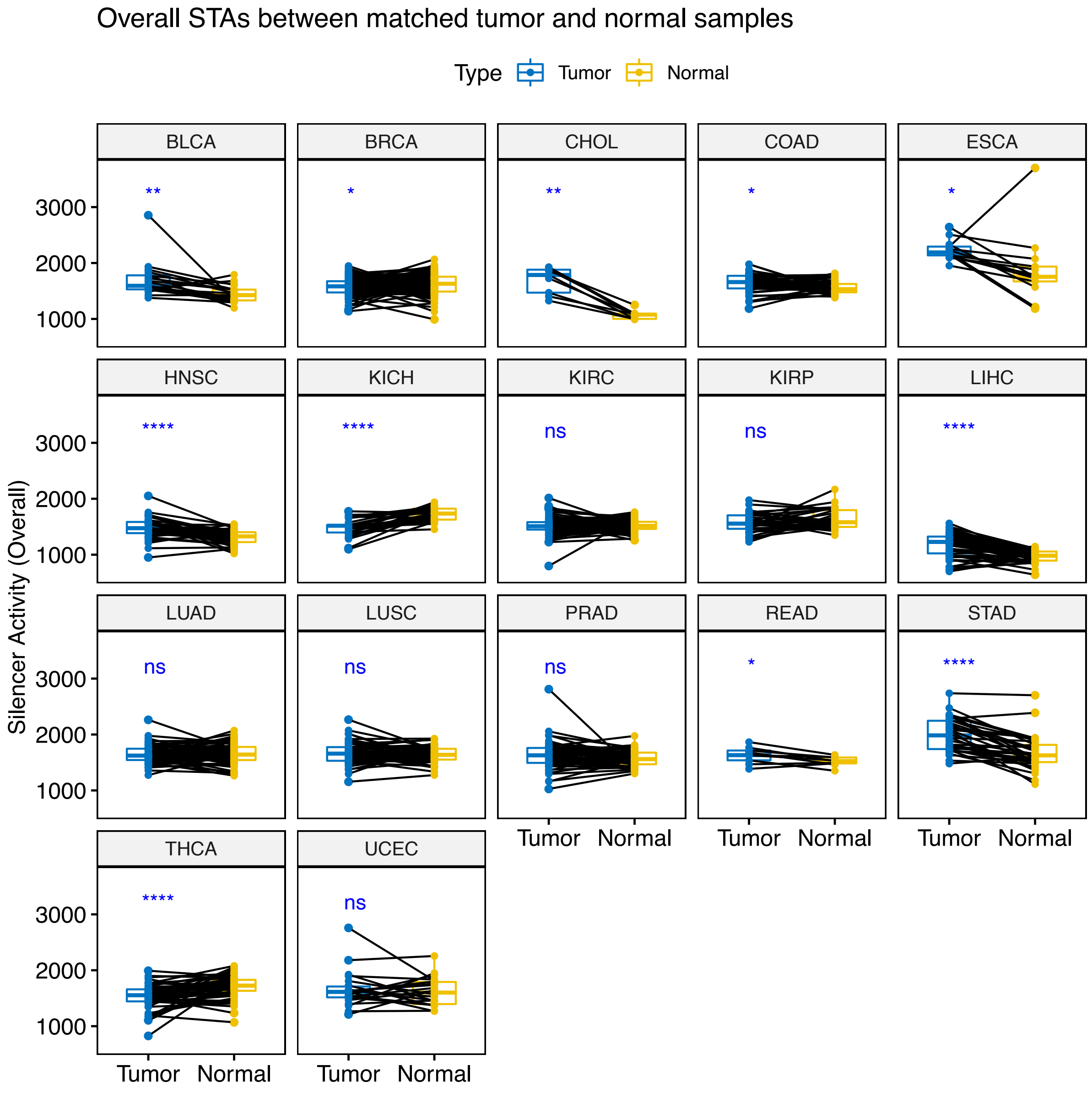

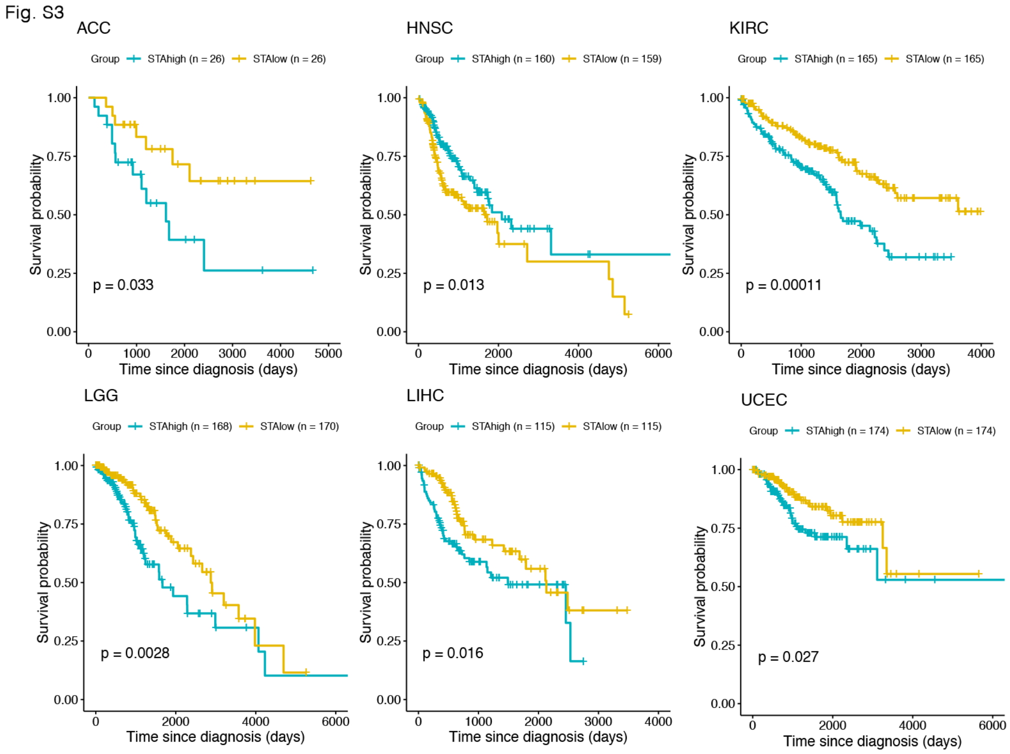

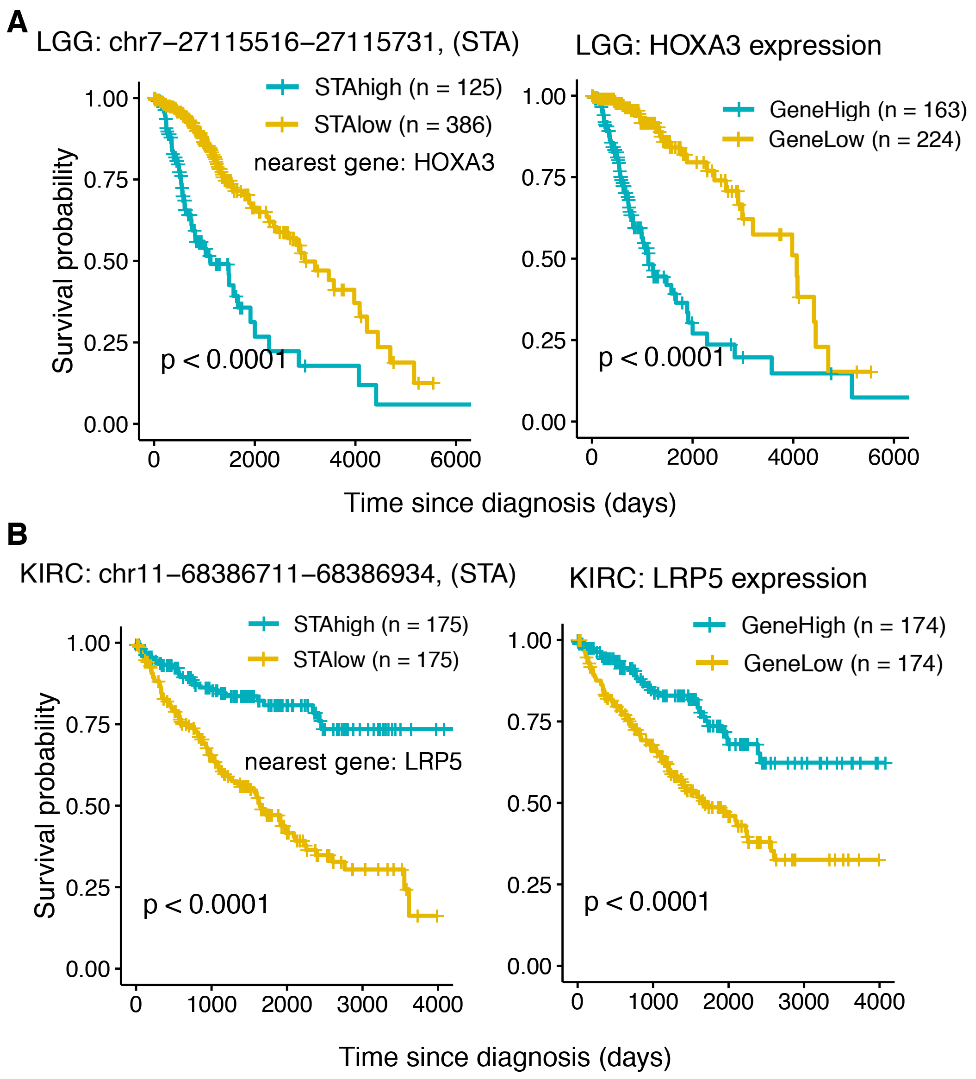

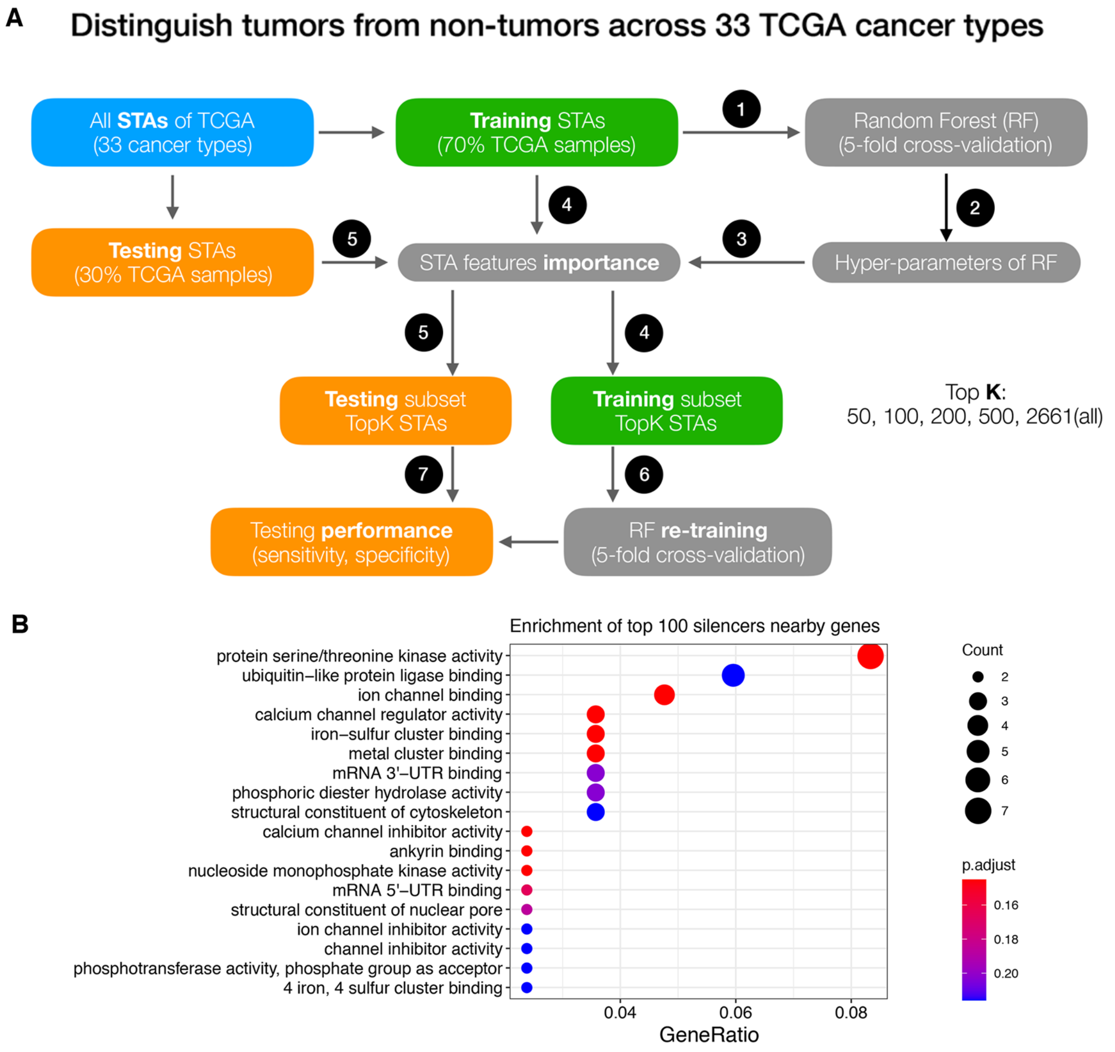

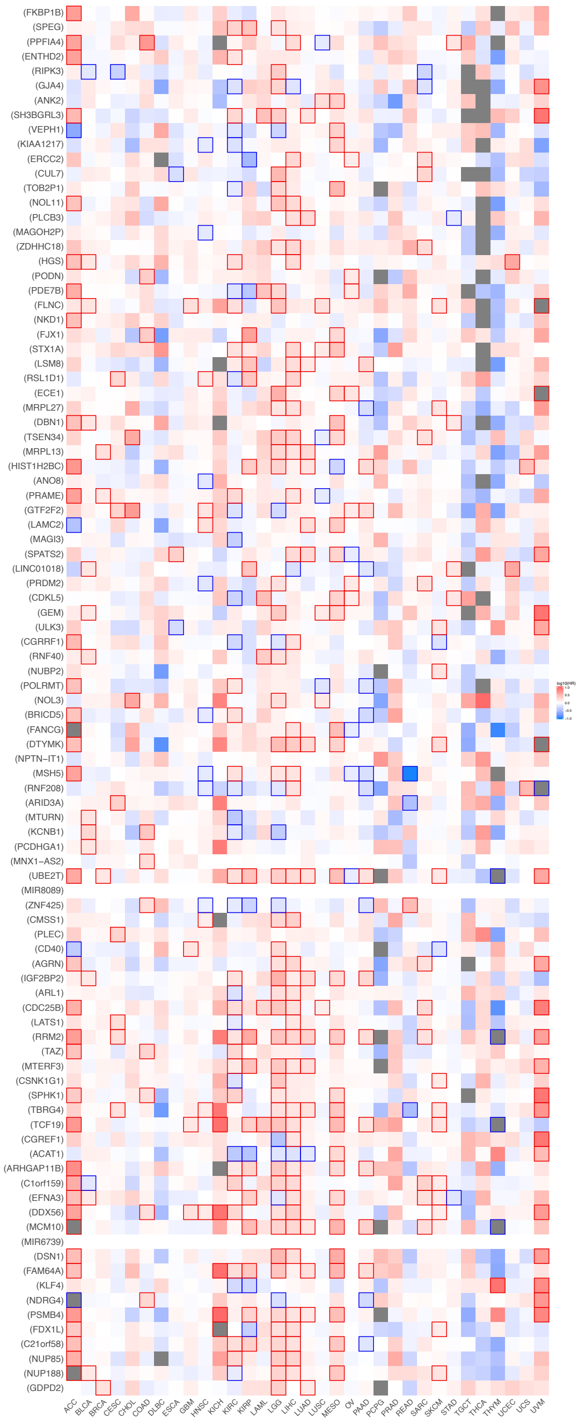

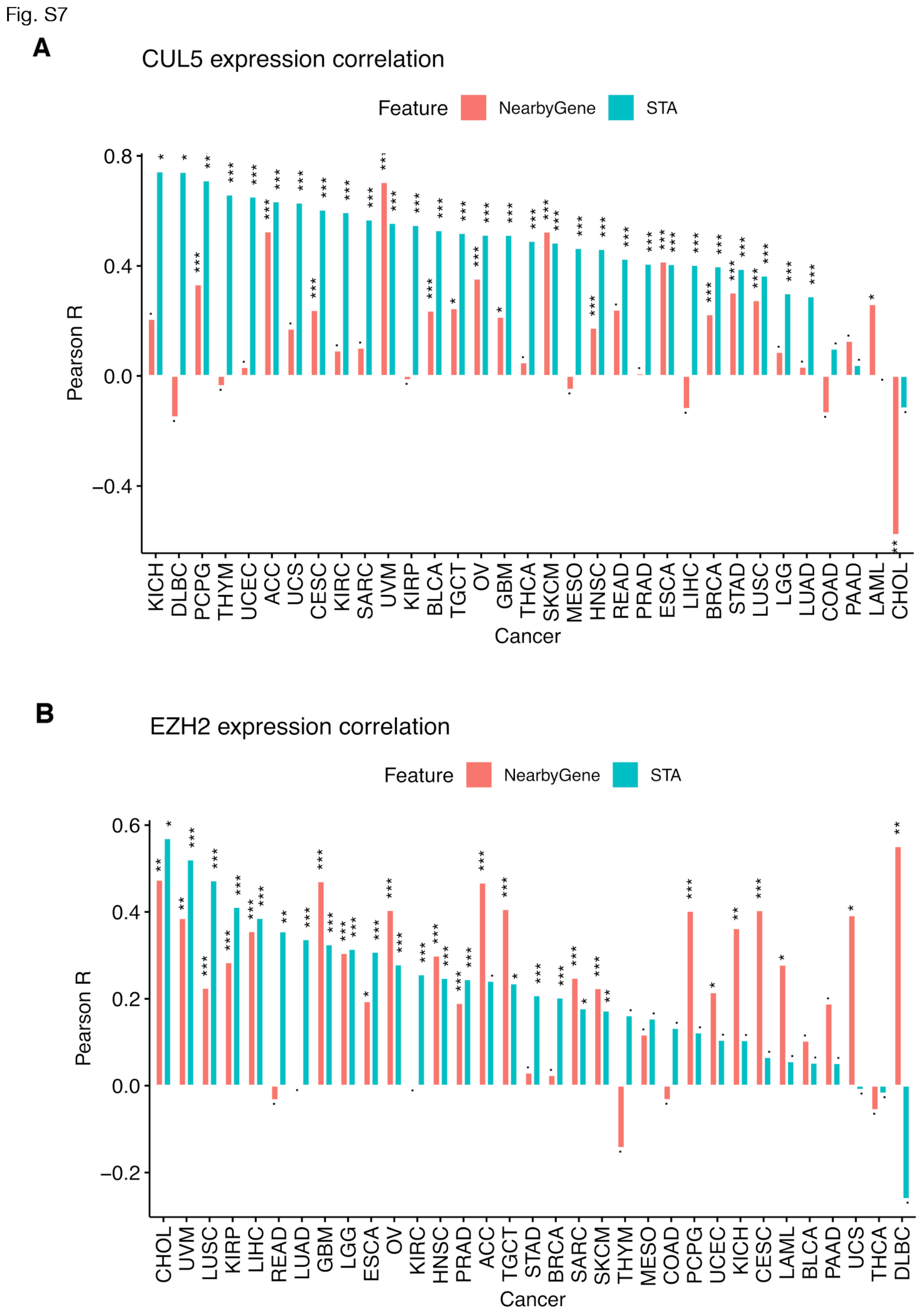

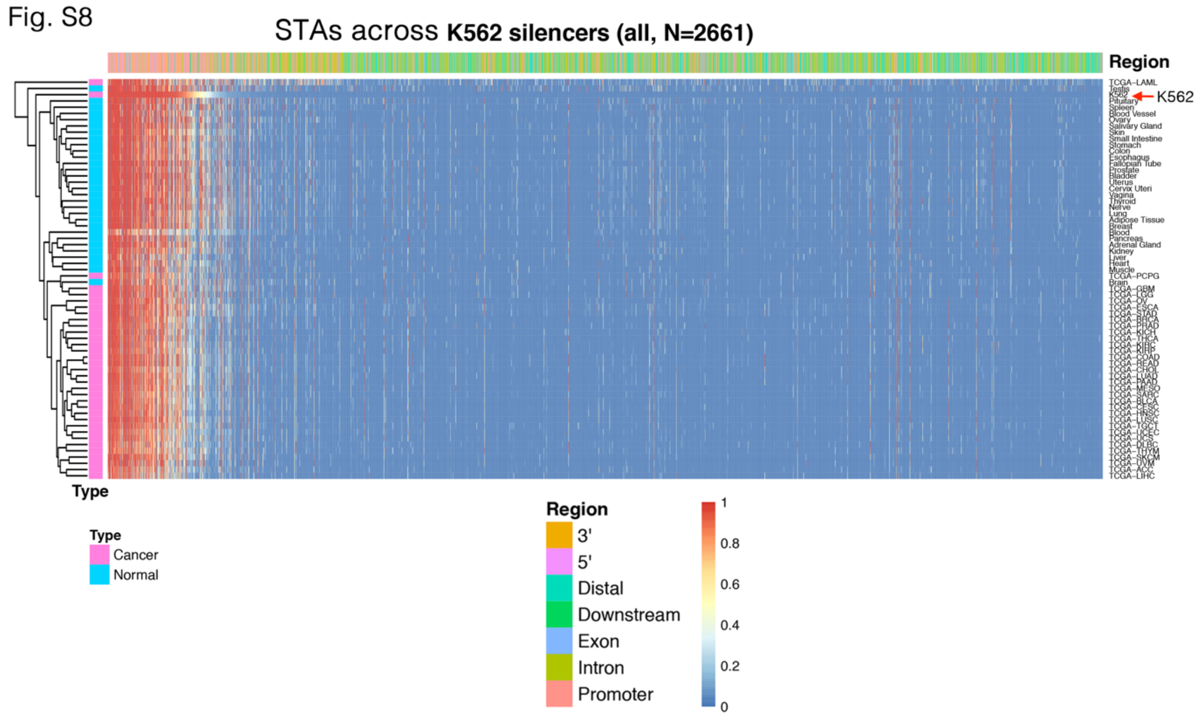

Fig. S9

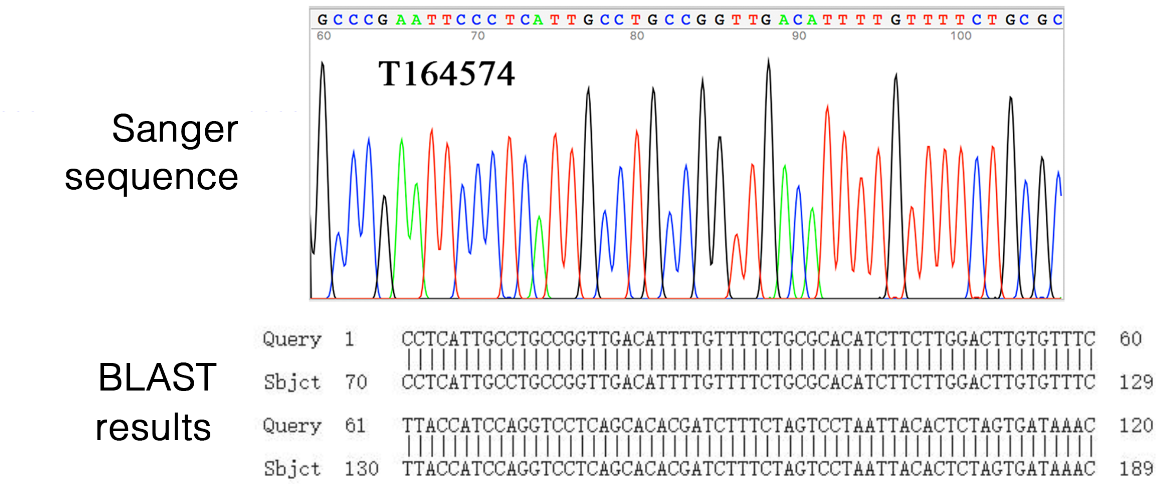


**Fig. S9.**

Fig. S10

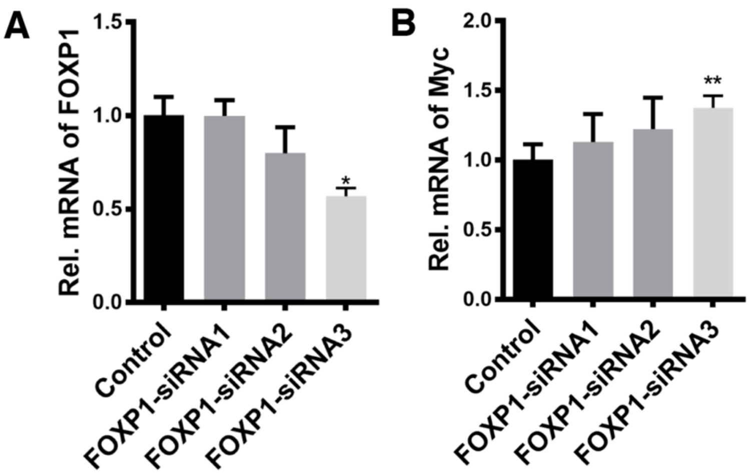


**Fig. S10.**

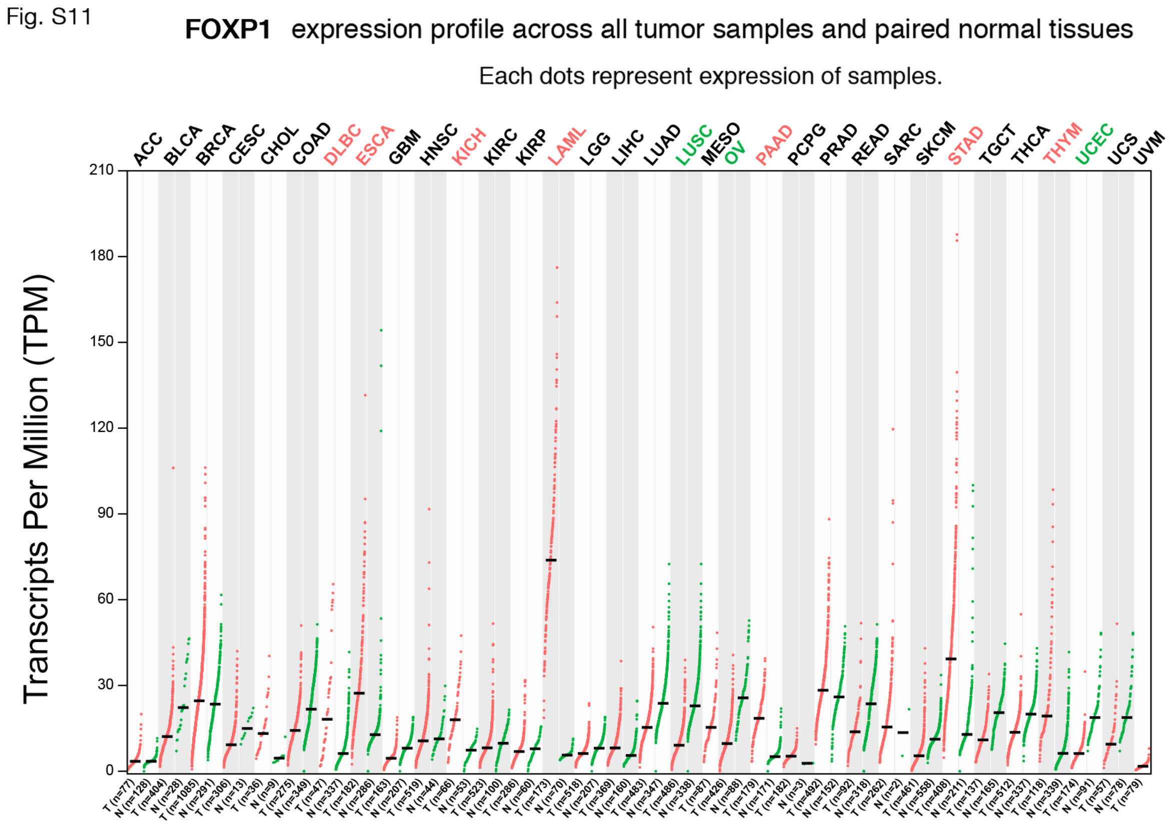
