## Supplemental File S1 for "Pan-cancer silencer transcription atlas reveals leukemia silencer hijacking FOXP1 to enhance *MYC* oncogene"

### FOXP1 binding motifs in slRNA T164574

ataattaatattatatattctagtaaattatatatataattttatatagccataatatgttgcattacaatgtatataaaatgtgtattatataacacttatatacatacagatgtatgtgtgtaaatacacatataGAGCC
CTTCATTATTTCCTCTGTCATTACCTTTTCACTTAAATTTTAAATCTTTAAACAATTTCTCCAGACTTCACACTCTGTAAAGCAAAGACCTGACATTTCCTACGTACACCGT
CATCTCTCCTCCCTAGATTCCACCCTTGACTTCCATCATATCTACTTCAGATTTATAATAAACTTTCGATTGTAGAACAGATTTAGATTTACTGAAAAACTTAGGTACTTC
AGAAAATTTACCCCTGAAAAATTGTGGGAAATTCTTTAGTCATTACTCTATCAAATAGAAGAGTGATTAAAAGTTCAAGCTCcagagtgactgatggtccaagctcggcaggcag
acatttagagttcaaatcatgcttttatgatttaagaacaccatgaccttggtaaagttacttgactttctgatgcctcagtctctttatacttaaaatgggaacaataatttcacaaatattatatgattataggcttataaa
tatttataaacattataggattattaaagagttaaaacatttataacacttagaccagtacctTTAAACTGGTGAATATGTATATATTAGACTTTGCTGGATTATTACCTATCTTTCAGTCTCTA
TTCTAACTTTCTGGGACATTGATTATTGTTATTGCACTGTTATCAATGTTGATtatgttaaaccttgctgtttttatcttttctatctcaatttatctttcattatttccttaccttgtccacctattattatg
agtaattttttttcctgatatatcttccagctcattgatttacttcttggctgatttgaatctgatgtcttaaacatccattatttttgttagtttgttttttatctgcaacatctgtacttgtaaaattgtagattaaataagttctaa
aacaaataaattatttaacaaataagattatttttcaaatctacttaaccactttttgaaaaaaatttttgagacaacatctcattctgtcacccacgctggagtacggtggcgcaatttgagctcattgcaacctctgcctc
ccaggctcaagctatcctccgcctcaaccttctgaagagctgagactacaggtgtgtgccaccatgcctggctagttaatgtattattgtagaggtggggtttccccatgttgcccaggctggtctcaaacttgtgaggtca
agtgatccgctcgcgttggcctcccaaaatgctgggattacaggcatgagccactgcacctagcctctacttaaccactttttctaatgcattatggcttgcttatctttaatcctatcttttatatcctgaattattatttttttt
gtaaactatttgggattattcagttccctgaagtacaagaaggtctaatttgttgttgccaatttgtccttattttctctggatttctgcttgccctcatatatgatggcatcatcctttggttattattattatttaccttaatcct
gtgtactgaattgaggcttcacttacacagataatatttacttattttttcaggcactgaaagagagtaacaacctggtattatctaaattgaaacaccctacttgggttcccataaccaaactgagatcatacatttgaag
tcaagacagaatcacagagatctatcattccagattctcaaaggatgctgttttttcttgttattactgaatttcctaaccagtcccatttccttctcagttcctttctctgcaagcagtcggtgttctaagctgtcacaagagat
gtgtcccatcaagagtttcTGAGAGCCTTTGCGTTGCCGTGGCCGCCAGCTGTGAACGCCGATCATAACGTCATGTTCCTATTTGCCATCAGTTGCAAACCTCAAATATTCTG
CCTTTTCATTTGTTCTCAGGAAAAGAATATACTCATCTCAAATAAAATCTTGAGGAAAGTGAAAAACTTGTACGTAAAGCTGGAAAAGACCAGGGAGAAGTGAAAAG
TCGTTCTGGAAAAGCGGTTCTGGAGCCTGGTAGTGGCCTCCAGTGGTGTGCGGGACAGGGCGGTCAGCTTGGGAGGACAGGTGGTCAGGCGCTGAGGACTTCGA
GCTGCTtggtgtcgggctggtgccttgatgttggtcaagctgccaatattgactgtggaaatcggcacataccacagagaaggactgttttcccccagagaactaagtgttaaacattttcaccaacaaaccaGTAC
AGATTTGCAACAAGGGCTTAACCTCTCTGGGGTTCAAACACTACTTGTGAACAATAAGGTAAGAACAGGCTGCTGTGAGAACTGTGTCATTGCATGCAGACAACCTG
CAGGTGTAAGCTCACAGTAAATGTGCATTCTCTTTATTGGGAAAACCGTTAGGACATGGAGCTAGTTCTTGGGCTGACTCAGAAAACCATGTGTGAGGCGGCATGGC
CCCACCTTCCCTGCTGAGCCCCGGGTGTGTAAACATTTGTAGAATAATATGCCATCGTGGAGCTTTGCATTCTAAAATGATCTTCAAATGAATTTTTTTAACTTTCTGAA
ATGCTCAAGACTCAAAAAATGCTTATTCATTCACCACCCAAAAGATACAGGCTGACTCCACCCAGTGCATGTTCCAAAGCTTTGTCCAATGTGTCTTTCTGAATTTTTAT
CTGAATAACCTGAATCCTGTATTCTGCGAGCCCGTCTCCTGCCTCTCACAAATATCACTCTAATTTCTTTATATTTCATATTCTTTTATAATTCCTTATAAGCACAACAGC
AAGGGAATCGCAATAACTCTACATGAACACGGAGTTTGCGCAAGAAATTTTCCACGtatcgctttggtccctacaacagttctcaaaattgcgtattatatgtccacatgcactaactgaagaa
aggaatttcagagagattcaattgcccgcattcatgtacaactaataaatgaatgtgctgggattcaaacctagccccttcaaatctccaaagtccgcgttcctaccgacccacctttcctccatctgCCTGCCCGTCC
ATCAGTGCCCACCCGATCACCTGCTCTCAGGACACCATGAGTGGTCTCCACTGAAACACTTAGAGTCTCCCCGTGCCTTATTTCTTCAGAAGTGACCCAACGGTGCAG
CTGCTGGCTCAGGAGAGGAAAGTGGATCTGGCCTGTCTGTCCTGATTTGTGTCTctctgaggccctgggcacagggctctgggctgtggctgatgggcAGAGGCCCTGGAGGACGCC
CAAGCTTGGTCACTGGTACTAGTGACTTAGAGGCTGCGTCCTTTGAGTTCAGAGTCTCAGTACGAGGCAGCCTTCCCTGGACACACATAGGAGCCCCTCTCTCTGGTC
AGCGTCATCGGTGCAGCTGATGCTGCTCTGGGCTTCCTCACAGTGACTTCAGCAATCCTAGGACACATGTTAGCACCCACCAGTAGCACGCTTGCCCGAAACGTCTG
GAGAACTTAGTTTCCACAAACCATCCATTTTCTGGTTTCATGTATTGTTTTCCCAGTGAAAATCAATGCTCAGTTTCCAGATCTGTGATCCCTGGGTCTCCCATGATCAC
ATTGGCCTCTGTTTCCTTCCTGGCACAGTGCTTCCGTTACCATGGTTGTCCAAGATGGATCGGGTTAAACATTAGCTGCATCCACATTTCATGTGGAGTCATCTAAAAT
GCCTTTCAATTCTCAGCTGTTTTCTTGTGTTAGGCATCTTTTTTAACTGTTCAGGTCACCCTTTCACTCCAAGTAGCTACAGCCAAAAGTTAACTTTGTATCATGTCCATA
TTCCTATACAGACTAATTTTTTCACAAAGCCTGATTTTCAAAATAGTAGTCAGACAAAAGACATGGATATCCAGAAAACAGACTCTTAGCCTCTAATTTCCATTTGTTTT
CTAAGCACTTGGATAAAAACTGATGAGCTCCCCACCAGGCacagcagagaagagtgcagagagaatcctgacctcatgcagcttgcattctgctgggagtaaggacagtgaaccaacacagaaat
aaaaactcagccagatgtcaagtaatggtgagttttctgaagaaaaatacagccaggaagagagaagttgtctctcagaggacagctttagccagatattccttaatcatcctctctggagagatgacatttgagaagg
ctgtggatcaaacagaggaatgaattagatgaagacacagagaaagaactttgacttgaggcagcaacgagcttggctgctggagaaaagcaagaaggccacatggcctgctgggaaaagttccaggaagaagct
cagccctaggagaatacaaaccgattggatcaggagacccctgagttggagatgaacattggtaaactcccctcattgccgtggtgaactccccTCGTCGCCGTGGTGAACTCCCCTCACTATCGTGGT
GAACTCCCCTCATCactgtggtgaactcccctcattaccattctaaaaacctggcccagggaagagcttctctgccatgttctacacattcgatgcatgaagaggcatgatcggtgatggcctcccctgcctccgctc
cacctctgcacgtggtgactcagctcacaagcccaggagaagccctgtgtcaccgtgttcgggaggcgctgttttgggagctctccctggtgttctcctgacttgttacaagtgatacaatccccttgttaaagcctccttg
gttgtggtgactggcttgtcgctcgccaagctatcgaacccacctgttgtgtgggcaacaGAAATAATTATCAGGACCCGTGTGGGTTGTACTAAACAGAAGCGCAGCCAGAAATGATGAA
CCTATGGCACCCACCACTACTTCTCCTCTCCATGtgtggtgcttttaaaagaagtccacacgtttcttgacatccctctgcccatcaactgctggggagtgtcctccccttgaatatgggctgaacgtaggca
ttggaccgccattggcagatggtgtaaatacgtgatgtgacttccctggcaagctcatcaaaggcaagtgcaggctctacctggtttgctggaatgctggcgccggagcctgagttggtgagaggtctcacccacctgag
cctccgtgctgcgagaaaatcAAGATTAGCTATGCAGAGTAGACGCATTGAGAGGCATCAGGGCCTCAAGAAAAGAGTGAGGCTGGCCGGTCCCATGTGCCTGAGAGCTC
TCTGTCTGCTCCTGCTACCATCCAGCCGCATGAGAGACCTTCTGCAAGGACAGCACAGTGAAGGCATTCCTACATTTTCTATTCTTAGACACCACAAGGGATAATAAT
CGACACTATTTTAAGCCACTATATTTTTTGGCAGCCACAGATCATTAGATTCCATTAATTCACTGACACTGGGCCCCTGTGGCTGCTGTGTCTCCCAGTGTCCCTCAACA
CCTGACCAAGCAGCCATTGTCAACTCTCTGGAGTTACTCAGGGCAAAGCTTTTCAACCATCCCTAAATGAAGCTGTTATGGAGGTGGGTCTGGGGTTGCAGACTGTG
GAAGTCTGTGGCTTCGCTGGGGTGCACAGGTAAAGACCACAGTCACCGGGGTCTTAGCTGATGAGAGCAGAGCACACAATGGTCAATGCCCCAATCCCCAGCAGCC
AGGTgcacctgcccagcacccgcccagcacccacccagcacccgcccagcacccgcccagcacccacccagcaccTGGAAGGCTTCAAACAGGTTCCAACATTTCTATTGTGTttggtatttcata
gtatgataattgataattgccatatttgacaattcgtattaagaattTCTggctgggtctggtggctcatgcccgtaatcccagcactttgggagtccgaggcaggtggatcacctgaggtcaggagttcgagaccagcc
tggccaacatggagaaaccctgtctctactaaagatacaaaaattagccgggcatggtggtgcacacctgggtgtaatcccagctactagaggggagctgaggcaggagaattgcttgaacccgggaggtggaggtt
gcagtgagccgagattgtgccaccatactccagcctgggcaacaagagagagactctgtctcaaaaaaaaaaaaaTTCTACGTGTAACATGCCATTAAAGAGTTTTGAAACCATTGAGTTATA
GCAAAACTCACATGCACACACACACACGCTCTACTTTAAAAGGCAAAGACCAACAGCTGTTTTTTTAGCCCTACGTGTCTCACTCCGTGATAATCGCAACTCTACATGC
TCAATTATTAGAGGTCTTTAAAATATGTAATTGGCTGACAGAGGTGAACTAGCTACACACATATGGAATAATCGTTTGGAATTCATCAACAATAAAATAATAAAAACC
GAAGACATTTCCAATTTTATAAAGACAGATAATTGTCAAAAGTAAAAGAAGGAAAATAACAAACAAACTTCTAAAGTCTATGCTCAGTTGGTGTTACCAACCAGCAG
TGTGCATTTCTGATATGGAAtttgttttgttttgtttgttttgctttTTAGTCTCAATCAATTATAAATCCCAAAGGTAAAAAACTGTTCTGAATTTGCTCCAAACCATCACTGCTGTC
CACTGAAGAAGTTAAAGCAGAACAAATTGTTATTAGATACAAGTGAGGAAAATTCCTGGAAGAGTCCTTACAATTTTAGGGATACCTATTAAAAAGGAAGAGAATG
ACATTGACTAGCTGAAGCCAGAGGTGCGGCAGTTTTATTTTGGAAAGAGTAGGTTGTCTTTAAATATGAAAGCTTAATGGTATGCAGCCTGTCGTCTGCTATATGAA
AAAGCCCATCAAGATGATGGGAATGTGTCTTAATAGGACTTGATAGAAGTATACAAAGCGGTTCCACATTTTGGTCACTCCACAGAAAGACATGCAGTTTAGTTCAT
CTTAAAGCAGTTAATGTTCTTAATAATTGCTTTGTTTTAGACTGCCTTTATGCGTTTGCTCAACACAATTGTTCTGTGCATACAGAGAGACGGGGGCATTTCAGAGCGT
TACTTACAATTATTAATAAACTATTTGGCTACTGAATATTCACACTGTGTAGCTTTTCAACTCTGCATACAAGGGCTGTCTATTGAAACTGCCACGGAGCCCCGGTGCA
CAGCTGGAGAGGAAAGCAGGCTTTTACACACCGGCCAGATCAATGCCCTGACAAAGTGGGAGAAGAATTGGGAGAGTGCTCTTGATATGTTGCCCTCTGTATACTG
GGAGGCAGGGTGTCCACCAGGACGTATTGATCAGCCGTGCCGTATGGATTCAAAGGTCTTCAGCGGCGCTTATGATATTTTACACCACATTAACTGTCGAGATGGTG
TGCTACGGCAGCCCTTTGCAAAACACAGGAGGGCCTCAAATAGCTGCAATTAAGCCCTCCAGAGAGCAGGTGCCCCCACCCGGGAGCATACAGTATTCCAGGGAAG
CGGGGAGAGGCCAAGAAAACCAATACTCGTCAGAGCTGCGGAGCGATCCTGCCCATCGGTTCCAGGCTGTCAGCACTTTCCTCCGCCGCAGCTTTTCATAATCCTCA
GCATATGCTCTTTGTTTTGATAGGCATGAAATTACATCAAGGTCATTTTTAATAACTGTCAATAGCTTGAGAGCAGCCGTCAGTTTTATTGACACCTACGCTGAAGAGC
TCAGGGAGCTGCAAGGAGCACGTCGCCCCTGCCTGGGAGCACCGATCCACTGGGAAACGGACGATTATTGATATTAGTAAATTAGTATTGATAGAAATTTGATGAG
TAGAGTGCATCTGGGTCACTCCAGAACGTGCGAATGTCAACATCCCAGAAAGAGCCGAGGCCCAGAGCTGTAAACAGACGTGCATTTATTGTCTACGCGGGGTTCCT
TTATCATTCTGTTCTTCTTCTTAAGAGACAAGCCTTTGTTTTAAATACTGGTCACTTACTCCCACTCCAAAAACCCAAAGCAGTGCTAAATACCCGCCCTTTAAATGTTAT
TTTCTTCTGTTTGGAGAAGAAATATTCCTTCCTCTGTACTTTATATTCTAATAGATCTCTGGAATTTTAATATATTTGCAAAGAAGATATTTGAATGGGACCTTATTTATG
GAATAGGTCGTCTCTTTTACACGAAGGCCTGATGGACCCCGAAGGGTCCCCAGTGGTTTTCTGGGGACGCTCAGGCTTGGCTTGCCCAGGGGATGCTGCGGAGGTC
TAGAAATCCTTTATAGATCAGGGAGGTCTCCGTATCTCAGCCCACAGACAGGCATGTTGCTTTCCAGAGTGAGCCGACGCAAGTTTCCATGGAAAATGTTGGAAAAA
GGGAAGACGGAGTTTGTCAGAACAGCAGCAAGCAGCATTGCTGTGTGGTGAGCTCGTGAAGCTGTGTCCTTAGAGGAAACTCACACCAGTAAGAGAAGACAACTG
TGCTGGCTTCTCACATAGGAGAGGCTGCAGATCCAGAGAAGAGAGGGATCGTGAGGAAAATTAAACTAGATACCCGGGAAAAATGAAGTCATATCGgaaggcagacat
aaagtcacttacgagttcatttacacaaaacgtccagaatacgcagctccacggtgacagcgagcaggttagtggagggctggactgggggacggggatgggatgactgcctactgggaaaagggcttattttgggg

76 agatggaaattccctagaattagattgtggcaatggctgcacagctgcgaatatatcaaaaacaccgaattgtacacattgaatgggtaaattctgtgatacatgagttatagctcaattaaagAAAATATAAACC
77 TGGAGAAGTCCCTGGGATAGCAAAAGTGAAAATCGTGAAATGGGAGTTAAATCTATTTATTTTAATTTAAGGTACAGATTTAGTAAATCAGGGAGTTAGTGTTCCAA
78 CCATTATGCCTTGTATTTCTAATTCTGTAAATTCAGAGCAAGATAAGATAACTAGGTGAAAAAATGCCTGTAGTACCAGGCCACTCTCAAGAAGGCGGGCTAACGAG
79 AAGAGGTGCTTTCTGTTCACACGAGGAAGCCGGCACGTGGAATGACTGCAGAGTTGCACAACCGGGCTGGGATTGCACGGGCCTCCTGCTCCTGTACAGCTCGGGT
80 ATATTTTAAACAGGTGGTCAATGAGCGACCAGGTTCACAGTGGCATTTTGAATTCTAAAACTTTTATTTCTACCAATCCCCGCGAGTCGGGTAACACGTTTATTGACCG
81 CAGGTGGCTGCTACGGGGGCCTCATTGCCTGCCGGTTGACATTTTGTTTTCTGCGCACATCTTCTTGGACTTGTGTTTCTTACCATCCAGGTCCTCAGCACACGATCTTT
82 CTAGTCCTAATTACACTCTAGTGATAAACCTACCCATAAACTCAGTGGCCCAAATGGTTTAACATGGGGTTATTTAGAAACATCTCTTTTAAAATATTTTCCCTCTTTGG
83 AAAATGTACAAAGTAGATAAAATTCTACCTCTGAGGGCTGTGTCTCACTTAGGAGTCTCCACTGGCTCATGGTCTCAGGAAGTGTCCCCGGATACTATGACAGGCGA
84 GATCTTCGCGTGCCCAGAGGGAGTCACAAGAGCATGGAGTGAGAGCAGAGCTTGGCGATGGTCATGGCTAACGTCTCCTGTTTACAAGTGAGGAGGCAGAGACCC
85 CAAGGCAAAGACAAGCGACTTATTTGTGGAATAGAATGTGCAAATCCAGATTTTTTAAGATAGAGAAACTAGATCATAAGAATGTTCAGACTGGAGGAAAAAATGC
86 AGATCATGCGTTCTGATGAGAAAAAGAAAAAACTAGCTTTGCCCTAGGTTCTTTCTGTGGTCCACTGAATGACACAGCTTCCAGCCTCCTGATTCTGTACTATCTTTCA
87 AATGCCAGACACCAAAAAAATCATAATTTATAGTATGACTTCACCTGAAGAGGGAATCCAATGAAGAGAAGTTACCGTGTAAAAACACAGAGTTTACATCAAGGTAC
88 TACAGGTTTTTATACAAGTATTGAATAAACAAAATGTATGCAACAGTGTGGAGGTGAATATGAACTCTCAGTGAAAATTTTCTAAACCCTTAAGAGGATTATCAGTCA
89 AGAAACAACATTCTATGATTCATCACGGTCTTCAGATGTGACCAACAGCATTGTCTCAAACATTTCTTCCTGCAGACTGGAGGTTAAACATCATCAGAGAGAGAATTT
90 ATCGTGAGGATTTTACTGGTTCTTTGTCAGAGGTACTTTCTGAGAAGGTATTTTGAGAAGCATACATTCAACTTAGTGAAGATGACTCTGATGTTTCTTGAAATTGTCA
91 CCAGCAGAATCTACTTACAACTCTGCTACCAACTGATTCAGAAAGTGTGTTTAAATGATTAGCAAAGAAAATCCCCATGTGATTACGTCCTAAGATTTTGGCCATTTTT
92 CCTAGATCTTCTAACAACTTTAGCACGGTCATAATTTATGACCAAAACAAAGGATGTAAAAAAAGATGACAAAAATAAAGAAAAATATAAACAAACTGGTGAGTAAA
93 CATCTAATACCAATTCTCTCAAATAGGTTCAATTGTATGGAATATGAAAACAATCTGTTTTGGCACAGCATTTAGCATAATCCTCGTTAAAAAGAGAATATTTTGAATA
94 ATAAATTTCAAATTATTTCCTCATACATGTCCATATGCGTAATAAACGAGTATTTTAGTTCCTGTCTAATTATCTAGATTGGTGCAGAATTTTTTTTAAAAAAAAATATG
95 CCAGTTTTAATTGCAAAAGTGCTGCCGGCTTCAGATTTTAATTGCAATTCACAAATGGTTGGACTGGCAGTGCTCCATTACAAAGAAAAAATCATTTTAATAGCAGAA
96 GTACTGCCTCGCTGCAGATTTCAATTTCATTTTACAAGATACTGGGTCATTCAGCGTCATTACAAGGAATAAAATAGCTGGTGTGAACCTCCGAGGTGTTGGCTGGGT
97 CATGATACTGCCTCATGTGCTGCATGGAGGCTGCCTTGAACTTGGCTCTTAATATCATCCGGAAGAGCAAAACCCAGAGAAATCCTAAAATCCTTTCAAAGGGGCTCC
98 CGTCCAGCCGCACTTGGCATAAAGCAGCCTCTCCTGTCCCTTACTTGTTCTGTCTTCTCTATCAAGGACTCCTTCAGGGTGAGAGCGACTTCAGTGTTTCTAATCTGTA
99 GAATAAAAGAAACGCAAGCTGTTTTCATGGAGATATTTTCCCTTCAGATGTTCCCATTCATCTTACCCTAAGCTCCCATGCATGTTTTCAGTGTGAGGCCTAGCTTTTCT
100 CTGACAGGAGGCTATAAGCACACAGGTGAAAGCCGTCGTAGGATTCACCTTGTCTTTGCCCAGGTGAAGGTTCTGGGGGTGGGTCTCCCCAGGTGACAGAGCACCC
101 TCATGACACTGGTGTCCCTGTTGGTCAGATGTTCTATCGGTCACATCACAGACTGTGAATCAAGAACAGCAGAGATGGAGAGAAGCGACTAAAGATGCCTTTCCCCT
102 CTGTCACGCAGTAAGACTCTGAACTTGAGTCTCTAGCAGCTGAACGTGGAGCTAAGCTGTACATGGTTCTCCTCGGCTCATCATTTTCTCATGGGGGTGACCTGGAG
103 GACATCCTGTGCTCAGGGAGGCCACCCTCCCACACAGGTGCCGCTGCACATGCCCGGGAGGGCACCTGCTCCGTCCCCACTCCAGGGCTATTACACCCAGTGTGGGC
104 ACAGCTCTCCTCCAACAACCACTGGTGAAACCCAAAGTGTGCTCTGCTGATGGAAGTCCTGAAGTCAGATAAGCAAATTAAAAGGCCAATATAATCTATGATTTAAAT
105 ACATTCCAAACAGATTTAAAATATCTTCCTTGAGCTCTGAAAAATTACAATATCTAAAGCCCTTGATTACTTTTTAAAATCTCTTATTTTAGGATTGTATCGATTTTTCAC
106 TACACATATTTTTATTTTTACAGTTTGGTAAAAGTAAATTATAAACAAATAACATGTATTCCATTATACAATTCACatatatatgcatatatttatctctatataatgatatctgtatatat
107 aACTTAACAAAACATGGAGGTACTGAGTCCACACATATCACTGCTAAAATAGTTATAGGAATGTAATAACAATTTTTACTTTATAATGACAGTAAATTCATTTGACATT
108 GAAATATAATAATTTTTTATGTTTTTAAAATGAGGCAGAAGATAAAGGTTAACTTTCTTAGCGTGCTATTACCAAATTCTCTTTATATTTTAAATAAATGGAAATGTTTT
109 ATAACTCAGAAAACATGATTCTCTCATTCCCAGAGTATCACACAATTTGAAAATATATAAATATGTATAGAGATAAATGTGTGCAAACAGAAAGGCTCCGCCTGGCAT
110 TTCCAGTGCACTTTTGTCCTAACTACTCTAATATATCTCAGCACAAATGAAGCACCATCATTATCCATTATGTATGACAGTGATGTTACATTTGAAGTGTATTTTTCTTAA
111 ATTCATCTCAAGGCACTATTAAAACAGAGTTAACAGTTAAATGGTTGCTATTTTGTGCATACAGAATAAGAGAGAATTGCATTCCCCATCTATTTCTGCAAATATTTAT
112 TGTAAGACGCTGTGCTCATCCCAGAGCTTGATAGCCAGAATGTCCGACCAGAGGTCAGTGAGGGCAGAGACACCGTATGCCCCGGACAGCGCTGCAATCTCAGCTG
113 CCCACCTATGTGGGAACTCTCACGGGGACTAAGTGCCCCAAGTCAAAACACTTACGTCATGTTACACAAAACCCTAAAATTCCCCCATGCAGTGTTGATGGTGTCTCG
114 GATGTGGCATCTTTCAAGGTGTCTGAACCTGGATTTACAGCACAAGCACGATGGCTCCACCCAGAAGCACACAGCGGCCTCTGAAAGGCTCTTCCGCTGCTGATACC
115 ACTCCATAAACGTCTAACCTTCTGCCCAGGATGGACCTGACTCTGGGGAAAACGATCCATTGCGTTCTGAAGTGTACTGCGAACATTAAAAAGGAGAAGAAGACACA
116 TAAATTGTGAGAAAGCCATAGCCAGTAAATCTGAGCGTGATTTACTTGAAAAACAGGAACTAGAATGAAATGCATGCTAAGAGCTTCCCACTGGGATGGGCGTTAA
117 CAGGGCTCAGCAAGCAGAATACGCCCGGGAGAGTCACCAAATGTCCACGCTAAGGTCGCTCCTACGCAGGGAGGAGGCTGAAGCTGGATGAATCCAAGAgtgtgcgg
118 gtgcctgtgagtgtgcgggtgcctgtgtttgtgcgtgcctgtgtgtgtgttctgtgtgtgtgcgcgcttctgtgtttgtgtgcgcattagtgtgtgtatgccttgtgtgcgtgtatgtgtgtgcgtgcatgtgcctgtgtgtgtgc
119 atttgtgtgtgtgcctgtgtgtgtgtgtgtgtgtatgCGGGGGGGGATTTCCCAAGAATATTCACTGAATCTGATCGAAACACACTTACAGGCAGTTAGCTGCTCAGGGAGGGAAAT
120 GTTCATCAAAAGCAGCGTTCTGCCGTTATGCTCAGGCGCGGGCACGCACAGGTGCGTGTGTCATGCACATACCTACTTGCCAGAACCAAAGCAAACCCTGCTCACAT
121 CCCAAGTTTTGCCCCAGCTCCCGCCCGTGGCACCTGGACTCCCGGAGGCGTGGGTGCAGCTCAGGCCCTCGGGTGAGGCCTGCCCGGTCTCAGGAGTGCCCAGGCG
122 GCGCTGGGCAGCAGCGAGCAGGTGGTTTGTGACAGAGACCCGCCCAGCCAGCCTGAGGTTCCAGACGGCGACAGGGGCGCGGGGAGAAGGCCCTGCAGGGGAG
123 GCTGATCCTGCGTGGACGCGAGGCCGTGGAAGGGAGGTCCCCCAGCCTTTGCCGAGGCCAGGCCGAGTGTGGATGCCCCTGGTCCTGGCTGGCCATGGGTGGTGT
124 GAGGGGCGAGGAGCAAGGCGTGAGGGTGAACCCCCGACCCCACAACGCCGAGGCAGAACCCCCTGGCAGGGTTCACGCCAGAGAGCAGAAGCCACGCGCAGGG
125 TGGAGATGGAGAAGGCAGGAGACAAGGACGTCCtccccttccctccccttccaccttccacctgtccaTCCAGGCGTCCGTGGCCTCTGCAAGGCGGCCCCACACCAATGCACC
126 CAGAGCAGGTGGAAGAGGCGGCCCTTGGGGGTTAAGAACAATTCCCATGCGGTGTTTCTGCAGGCAGggggcaaagaaaatttgcagtttgaacatcctgcccgttcccactgtgcga
127 tgtggcacaacttgctcgattttgctgggcgttgattttcttatctggaaaatggggagccccccctcgggggccggtgtgagAGCCAGCATCCCAGTGATCCCAAGGGCCACCCATCAAGACCATTG
128 CGCAGGCTGGGAGGCGCCCCGGCCCCCTGCGTTCTGCGCACCCTCACACTGCAGAGCAAGGCCCTGGCTGGTGGGCCACCTCCCTGTCAGTTTGAAtccactagcatttcc
129 tatggagcttcctccatggccgtgttaggtctaatgaggtctacaaatttcactacaatatagtccttgccctcaaggaactttcagcctaCAAAGTAAGTGTCTCCCTCAGTTTCCCTGTGCACTAAAGT
130 GTTTAGACAACTATAACCATCCTCTGGTAGCAATTTGCAGAGTGTCACAGGTTACAAAGTGCTTTATAAGCGCTAATTATCACTTCAGTAAATTATTTTTACATTTTTTC
131 CCAAAAGAGTGAAATAAAACAGCATAAGCAGAAAAACATAAAACGCAAGAATTGTTTTATGGGCCCCTGGCTGAGATTTCTGAACTTGAACATCATACGCAGGGAA
132 ACATTAGATGCAGTAATTATATAATGATGTTTCACTGTATAACTATTATCTTTCTCATTGTCACTTCTTTCAGATGAGGGCTGAAGGGATTAACCCTGTCTATTCTGTGC
133 TCAGAACAGCTTTCTAGACCCGATGTTTCCCTTAAACTCATCCATGAGCATTCAGAATTTACATGACCTAAGAGAATCTCGCAGATCCTAGAATCACTTCTAAACTTTAT
134 CAATTATTCCTTGCTTTCTCAAAATATCAGGAATTCAAAGAAGGGTGCTAAAGCATCAAATTGAAATAGATGTAGGTAAAGGGACTTAGTGGGTATGTATTTAGTTAT
135 AGATGTATTTAAGATGGAATGAATAAATGAGTTAAATGAATTGTGCACGGATCTTGTGCTATTTCTAATTTTAATAGTACTTGAAATCAGCTATGCTTCATAGCGGCA
136 GCTCAGGAACAGGCGTGGCCTGGTCTGAAACGATCAAGCCAAGAAATGAAGGACCTGATTTCGCTGGAATGCCTGCTGGAACAATGGCACATAGTGCTGACCAGG
137 ACAGAGGCAGATGCAATAAACAAAGGTAAAAAGCTGTGGGTACTGTGGGTAAGGTAGGAGTTTGGGGAAAGATGCACATCTGCACCAGTGGGATTATCAGATACC
138 CAGGAGCAAATGAACAAGCTTGGTGTTTACAAACAGTAGCCCTGTTTGTAAACACCAGATAAAGTTTATTTAAATTCAGTTTAGAGAGAAATGCCAGTCCTTGGGAA
139 CCCTACTATGCAGAATAACACCAGCGATGAGATGTCTGCGCACATGGAGGACCTCCCCACTGTTCGCGGCTCTGGTGGTCTTACCTGGGTGCCTCTGTAATCCTAGGA
140 AATTACTGCTTCCCGGTGATGTGCTTGGAGAATACAAGCAGGCAGGTGTGTGTGCTCACTCTAACTTTATGGGTGTTAATAACTTACATAGATAACATTTGTGTACTC
141 AATTAAAATTGAGCATTACTGAACTTTtatatatatacatggtttttaaacttatatataggtaatgtatgtatactattatatatgtattatgtatagatacgtgtatatacaca
